# Plasticity of host cell organelles in response to Pb infection

**DOI:** 10.1101/2023.04.28.538786

**Authors:** Mohammad Djavaheri, Leah Clothier, Colin Kindrachuk, M. Hossein Borhan

**Affiliations:** Agriculture and Agri-Food Canada Saskatoon Research Centre, 107 Science Place, Saskatoon, SK S7N 0X2 Canada

## Abstract

*Plasmodiophora brassicae* (*Pb*) is the most important member of the plant pathogenic protist subgroup, Rhizaria. *Pb* is an obligate soil-borne plant root pathogen which makes the study of *Pb* biology and its interaction with its host challenging. To overcome this challenge, we adopted an axenic cell culture method to study the interaction of *Pb* with its host plant *Brassica spp*. at the cellular and molecular levels. *Pb*, during its life cycle within the host, was confined to the cytoplasm of the infected cells. Our results showed that upon Pb infection, the dynamics of the plant cell organelles including mitochondria, plastids, nuclei, vacuoles, vesicles were altered. Both Plant’s and parasite’s *diacylglycerol acyltransferases* (*DGAT*) genes were highly expressed in the calli, which indicated a need for the oil accumulation at the switch between replication and growth stages of the pathogen. This notion was supported by the accumulation of oil bodies that were only observed inside the pathogen cells. Our results also indicated that starch was utilized by the *Pb* in infected cells, an indication that *Pb* induces cell viability by altering the host carbon metabolism. The ongoing dynamic changes in the infected plant cell organelles suggest that plant responds to *Pb* infection, through the induction of organellar plasticity.

## Introduction

Protists are a group of unicellular eukaryotic organisms that includes some of the pathogens of human, animals and plants. Plant pathogens of the kingdom protist belong to the supergroup Rhizaria with *Plasmodiophora brassicae* (*Pb*), the causal agent of clubroot disease of *Brassica sp*. being the most economically important plant pathogen.

*Pb* is a soil-inhabitant obligate biotroph and requires a living host for the completion of its life cycle, thus making it very difficult to study. Amongst the Canadian *P*. *brassicae* pathotypes, *Pb3* is highly prevalent in Western Canada (Strelkov et al. 2018). Genomics of *P*. *brassicae* gained momentum after the release of the first draft genomes of the pathogen (Rolf et al. 2016; Schwelm et al. 2015). The small genome size of *Pb3* (Rolf et al. 2016) and features such as a multitude of genes involved in sugar transport (*SWEET*) and dependency on the host nutrients such as sugar together with a reduced number of CAZymes (Rolfe et al. 2016) are common characteristics among obligate biotrophic plant pathogens. Recent studies suggested a tight relationship between sugar translocation and gall formation in the *Pb*-infected tissues (Li et al. 2018; Walerowski et al. 2018). *Pb* infection may trigger active sugar translocation between the sugar producing (source) and infected (sink) tissues. For example, *AtSWEET11* and *AtSWEET12* sugar transporters play role in the local distribution of sugars toward the pathogen in the roots of *Arabidopsis thaliana* plants infected with *Pb*, and mutation in these genes result in slower disease progression (Walerowski et al. 2018).

Accumulation of oil droplet inside *Pb* zoospores and resting spores (RS) has been reported (Aist and Williams 1971; Bi et al. 2016). Based on transcriptional analyses, Schwelm et al. (2016) suggested that fatty acids are synthesized in the plasmodia and degraded in the resting spore. Oil bodies are accumulated inside most protist cells; and are suggested to have additional roles than just being lipid storage pool bodies (Barbosa et al. 2015). Lipids have been suggested to be the preferred growth substrate of the protist *Naegleria gruberi* (Bexkens et al. 2018).

One form of the fatty acids storage in lipid droplets is the energy-rich Triglyceride (TGA). TGAs are used in the cells to store unused energy for later use (Debnath et al. 2017; Bi et al. 2016). Furthermore, TGA is used in the formation of lipid bilayer membranes as well as providing thermal insulation for the cell. Triglycerides cannot pass through cell membranes freely; therefore, one could hypothesize that organelle generation inside the *Pb* cytoplasm including endoplasmic reticulum (ER) and membrane for the growing amoebae and resting spores (RS) relies on the ability of the parasite cells to induce accumulation of TGA sourced from their own or infected plant cells. Lipoprotein lipases break down triglycerides into free fatty acids and glycerol. Fatty acids can then be taken up by the cells via the fatty acid transporter (FAT). *Pb*3 has two triglycerol lipase genes with an N-terminal signal peptide (Rolfe et al. 2016) suggesting that these enzymes are potentially secreted into the host for the break-down and subsequent uptake of the plant-produced fatty acids.

Organelle dynamics in plants infected with pathogens have been described for biotrophic pathogens (An et al. 2006; Glingston et al. 2019; Schulze-Lefert 2004). An et al. (2006) described the involvement of the multivesicular bodies (MVBs) in the defence response of barley to powdery mildew fungus. MVBs were suggested to act as building blocks of the papilla and antimicrobial compounds like reactive oxygen species that are released at the site of fungal entry. The dynamic of secretory systems is suggested to be a conserved mechanism by which different plant species release antifungal compounds (Schulze-Lefert 2004). Moreover, the active movement of actin filaments has been shown to facilitate the transport of membrane organelles (Semenova et al. 2008).

To further investigate the above hypotheses and understand the interaction of *P*. *brassicae* and its host at the cellular level we optimised a highly pure (axenic) culture method of *Pb* and conducted microscopy as well as molecular studies. Live cell imaging by means of light microscopy (LM), epifluorescence imaging (EM), scanning electron microscopy (SEM) and transmission electron microscopy (TEM) helped us to gain insight into the *Pb* infection process and phenotypic changes during the infection of canola (*Brassica napus*) cells. Here we describe the plant root subcellular changes that occur inside the *Pb-*axenic cultures of canola.

Our data strongly suggest that, over time, starch is consumed by clubroot as a source of energy, while oil droplets accumulate within the infected cells as a marker for the healthy growth of an intracellular protist. Additionally, changes in the dynamics of intracellular plant organelles in response to *Pb* infection are discussed.

## Results

### Brassica axenic cell culture supports growth and proliferation of *Plasmodiophora brassicae*

Cross sections taken from galls at four weeks post-inoculation with single spore purified *Pb*3 and from uninfected (healthy) roots of *Brassica napus* cv. DH12075 (hereafter DH12075) were disinfected and placed on Murashig and Skoog (MS) media. In a few weeks, dark calli emerged from the vascular tissues (Suppl Figure 1). Infected tissues of DH12075 produced the dark calli faster and in a higher rate than the healthy uninfected root explants. After a month, the *Pb*3 infected dark calli were transferred to fresh media, where they subsequently changed into friable white fluffy (WF) calli. Additionally, the color of calli gradually changed from WF into brown cells which over time turned into black. The old brown infected calli were able to regenerate the white fluffy calli continuously (Suppl Figure 2). However, calli from roots of uninfected control plants generated only a very limited number of cells on MS media, which turned brown and failed to regenerate younger calli. At the same time, single spore isolates of all the previously characterised *Pb* pathotypes, i.e., *Pb*2, *Pb*3, *Pb*5, *Pb*5X, *Pb*6 and *Pb*8 were used to infect the common susceptible *Brassica rapa* ssp. *Pekinensis* cv. Granaat. Axenic cultures of cv. Granaat-infected roots produced similar calli as observed for DH12075 (Suppl Figure 2). In addition, all the pathotypes were able to reproduce younger calli in cv. Granaat.

Because Pb3 is the most prevalent and most studied pathotype in Canada (Rolf et al. 2016; Strelkov and Hwang 2014) subsequent studies to determine the intracellular infection stages were conducted using *Pb*3 calli generated on DH12075 . The *Pb3* calli were maintained and studied on MS media for over three years.

Dark field as well as SEM micrographs of the healthy calli were compared to those of infected white fluffy, brown and black calli (Figure 1). The size of *B*. *napus* cells was significantly increased in *Pb*-infected axenic cells compared to the cells of uninfected calli in the absence of exogenous plant growth regulators. Furthermore, the cell enlargement continued over time in the axenic cultures transitioning from WF –longitudinal growth- to black –hypertrophic growth (Figure 1). Inoculums prepared from *Pb3* axenic cultures were infectious on DH12075 and produced galls on DH12075 and Arabidopsis (Suppl Figure 1). Cumulatively, these data showed that Pb was able to complete its life cycle inside the infected cells of DH12075.

**Figure 1.**
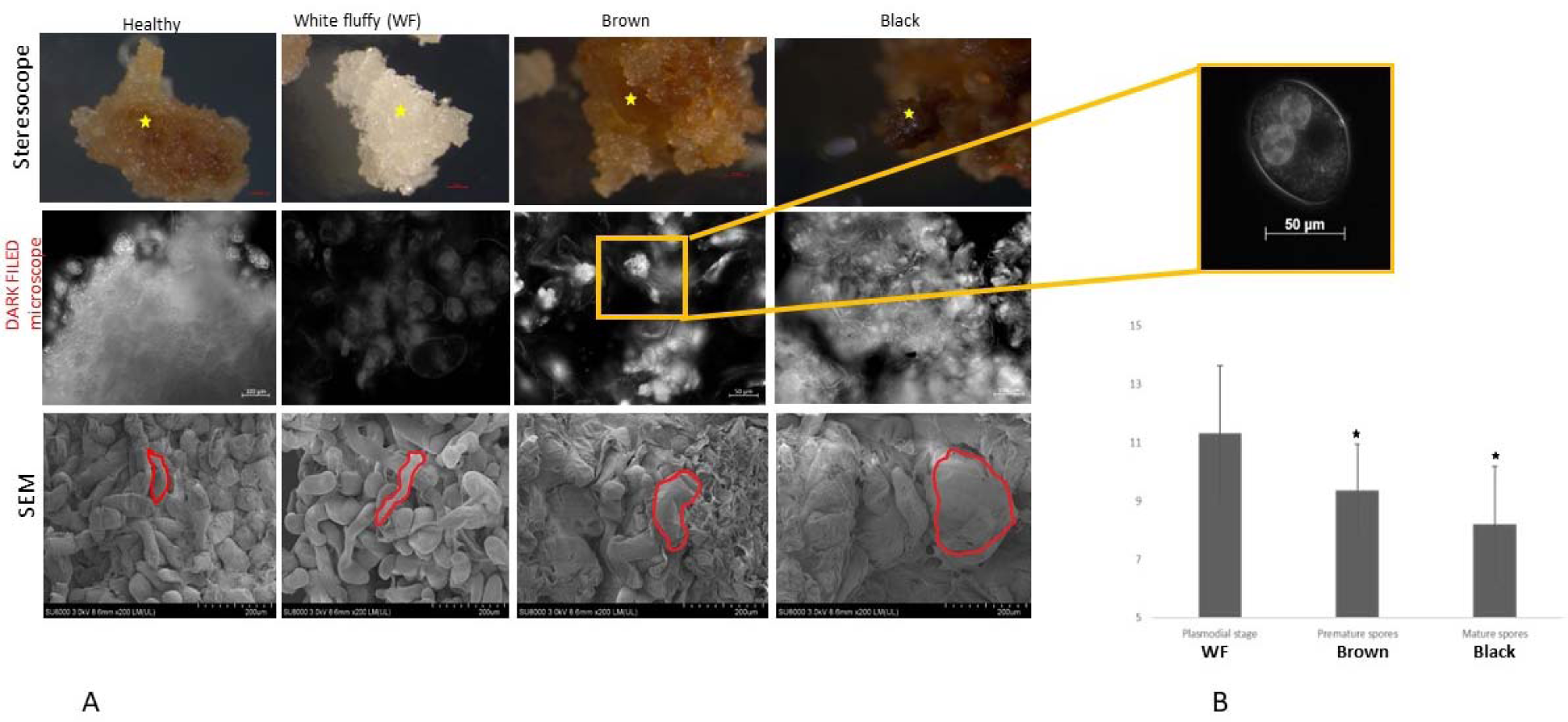
Different stages of calli regenerated from the *Pb*3 infected galls) compared with calli produced from healthy root tissues. A) Gradual discoloration of the calli cells in axenic cultures of Pb3 in *Brassica napus* cv. DH12075. The top row consists of the external phenotype of the different stages calli, under stereoscope; scale bars 100um. Axenic cell cultures include white fluffy (WF), brown and black calli from *Plasmodiophora brassicae* (*Pb*)-infected cells. The middle row shows calli under the light microscope (LM) with Dark Field. There are internal differences between the cells in various stages of the pathogen growth, in comparison with the healthy control calli; scale bars 50 or 100 μm. The square box shows an enlarged view of a brassica cell containing active swimming cytoplasmic zoospores as well as two pathogen-derived zoosporangia within the host cytoplasm: scale bar 50 μm. The bottom row is a representation of the cells under scanning electron microscope (SEM); scale bars 200 μm. Cell longitudinal enlargement and hypertrophy could be observed as a result of the infection by *Pb*. B) The expression of Pb Benzoic acid/salicylic acid methyl transferase gene of (*PbBSMT*) was compared to that of *BnActin* in infected calli. Bars are the averages of the expression of the *PbBSMT* (6 replicates) in WF, Brown and Black cells using droplet digital PCR (ddPCR). Asterisks show the significant differences of the values (Student t-test *P* ≤ 0.05) between Brown and black cells compared to WF.

DNA sequencing of the one-year-old axenic cultures of Pb3 using illumina Miseq, confirmed the presence of the pathogen within the cultured tissues with over 11% of the total sequencing reads being from Pb3 covering the entire Pb genome (Suppl Table 1).

*Pb* secreted effector *PbBSMT* has been shown to effectively suppress SA-dependent responses in clubroot-infected plants (Djavaheri et al. 2018). We measured the expression of *PbBSMT* in various stages of axenic cultures, i.e., WF, brown and black calli. *PbBSMT* was highly expressed in all three stages of the *Pb3* cultures (Figure 1B) and showed a similar expression profile as reported during natural infection of *B*. *napus* root (Djavaheri et al. 2018), further supporting that Pb growth in the axenic culture shares the same characteristic as the *Pb*3 growth in naturally-infected brassica roots.

### *Pb*-infected cell culture displayed infection stages similar to those in infected plant roots

Transmission and scanning electron micrographs (TEM and SEM) revealed that Pb3 was able to complete its life cycle and generate all the major infectious stages including zoospores, plasmodia, and resting spores within the axenic cell cultures and that the infection by Pb3 induced the regeneration of new WF cells from late brown cells.

Infectivity of spores obtained from the axenic cultures (Suppl Figure 1), abundance of pathogen mass (based on DNA sequencing; Suppl Table 1) and similarity of the gene expression profile of *Pb* effectors with the gene expression profile of Pb-infected plant roots indicated that calli generated from infected galls are appropriate tissue for cellular study of *Pb* life cycle and infection. By conducting microscopy studies of all the stages of the axenic cultures we were able to identify *Pb*3 primary plasmodia, gametangia/ zoosrporangia/ parasitophorus vacuoles (n), and secondary plasmodia (2n) in the axenic cells (Figure 2; Suppl Figure 4, and 5). Secondary zoospores were found to be actively moving inside the cytoplasm of WF calli cells, suggesting that they were about to be discharged from zoosporangia (Suppl movie 1). We observed zoosporangia and primary plasmodia inside the WF cells (Suppl Figure 4). *Pb* zoosporangia were normally present in three or more whorled lobular vesicles inside the cells and resembled parasitophorous vacuoles in Toxoplasma (Clough and Frickel 2017).

In order to study the early infection stages, WF calli were incubated with zoospores overnight. SEM micrographs showed round to elongated encysted zoospores on the surface of the WF cells (Suppl. Figures 4, and 5).

Studying TEM micrographs revealed a massive group of the primary zoospores inside the cytoplasm of the newly infected cells oneday post-infection (Figure 2A; Suppl Figure 5). Moreover, we noticed that plasmogamy of the primary zoospores, presumed to be important for *Plasmodiophora* to develop into secondary plasmodial stage, occurred either outside or inside the infected WF plant cells (Suppl Figure 4).

Brown calli mainly contained large and well-grown secondary plasmodia ready to develop into resting spores (RS) as well as some ring-form stages of the pathogen (Suppl Figure 7) resembling *Plasmodium falciparum* – the causal agent of malaria in human-inside infected erythrocytes (Tilley et al. 2011). In this amoebae-like growth phase, secondary plasmodia could be observed in the cytoplasm. The plasmodia varied in size and number. Black calli contained RS in various stages of induction within plasmodia (Suppl Figure 4). This coincided with the formation of resting spores from secondary plasmodia.

We noticed cell wall constraints in WF, brown and black calli indicating a prerequisite for an expanding cell (Suppl Figure 3). Interestingly, the plant cell wall thickness was drastically increased in black calli compared to younger stages of the axenic cultures (Suppl Figure 3).

### Starch metabolism in infected single cells

Cells of uninfected control calli were filled with large globular granules, contrary to the infected axenic cells that had few granules, as revealed by SEM (Figure 3). Tissues were stained with potassium iodine (Lugol) that binds to starch, noticeable as blue color. Staining proved the presence of starch granules in the healthy cells which were more abundant and larger in size compared to the infected calli (Figure 3). Additionally, starch was significantly less accumulated in black calli (LM diameter: 0.1-1 µm), compared to WF (2-25 µm diameter), and brown calli (LM: 1-2 µm diameter), respectively. Similarly, a reduction in starch granules in the infected cells was also observed (Figure 3), suggesting that starch is utilized in the infected cells. To find out whether the starch was utilized by the pathogen or the plant cell, we measured the expression of the *Pb* and DH12075 genes with amylase activity in infected calli cells. *Pb*3 has a hypothetical secreted protein with alpha amylase activity (PbPT3Sc00048_S_4.297_1), and a protein (PbPT3Sc00094_A_0.228_1) with homology to a protein involved in hydrolyzing gelatinized starch in *Rhizopus*. Expression of *Pb amylase* genes was compared to two candidate amylase genes of *B*. *napus* (alpha-amylase 1; GenBank NC_027757.2; and beta-amylase 8 ; GenBank NC_027768.2). Alpha amylase breaks the starch glucose chain at random points. In contrast, beta amylase acts on the bond between the last 2 and 3 glucose residues in the starch chain, releasing the disaccharide sugar molecules.

**Figure 2.**
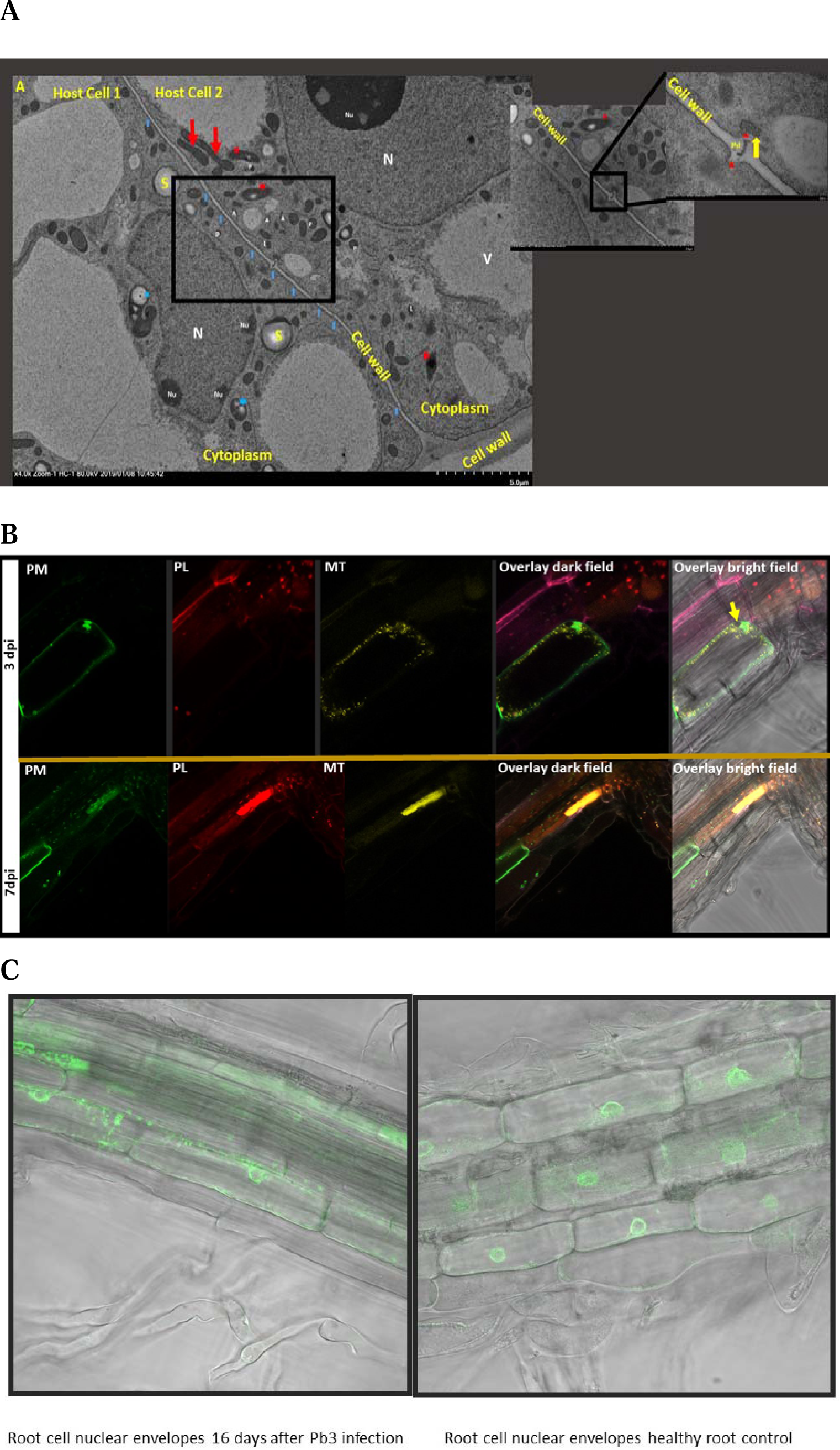
Changes in the dynamic of the plant cell organelles in *Pb*-infected cells. A) Transmission Electron Micrograph (TEM) of the calli cells 12 hrs after infection with *Pb3* single spores. Cells cytoplasm are filled with infectious *Pb* zoospores (red asterisks), and swollen zoospores transforming into plasmodia (blue asterisks). Infection by *Pb* causes drastic changes in the dynamic of the host cell organelles such as accumulation of plant cell mitochondria (red arrows), enlargement of the host nucleus (N), partitioning/fragmentation of the host nucleoli (Nu), as well as accumulation of actin filaments, and host plastids around plasmodesmata (blue arrowhead). *Pb* at the early stages of the infection tends to accumulate near the host plasmodesmata which is likely crucial for the intercellular movement of *Pb*. The square box is an enlarged view of a young plasmodium of *Pb* (yellow arrow) passing through the host cell Plasmodesmata to the neighbouring cell (Pd). Note that there are two papillae (red arrowhead) formed at the site of entry, where the *Pb* highjacks the excretion mechanism of the cell, thereby trespassing the neighbouring cell cytoplasms. A: actin filaments; V: Host main vacuole; L: host lysosome; S: Starch granule; P: Host plastid. From left to right, the scale bars of the pictures are 5 μm, 2 μm and 500 nm, respectively. B) Extensive dynamics inside the cytoplasm of *Pb*-infected cortical cells in Arabidopsis root at early infection i.e., 3 and 7 days post-inoculation (dpi). The top row shows changes at 3 dpi and the bottom row shows changes at 7 dpi. These pictures are generated using an Arabidopsis transgenic line tagged with cyan, red, yellow, and green fluorescent proteins (Kato et al., 2008). showing nuclei, plastids (PL), mitochondria (MT), and plasma membranes (PM), respectively. Pictures of Cyan-tagged nuclei were removed because the tag did not express well in the roots. At 3 dpi an inward extension of the host PM at the port of *Pb* zoospore entry and the shrinkage of the PM is observed in a root cortical cell. Moreover, PL and MT are accumulated inside the cytoplasm and their accumulation is significantly higher at the Pb penetration site. PL is also significantly accumulated in the cells adjacent to the Pb-infected cell. Overlay pictures show that PL and MT accumulation are co-localized. Disintegration of PM as well as strong localized accumulation of PL and MT is observed at 7dpi in the infected cell adjacent to the root vascular system. The PL and MT organelles are co-localized in the infected cell. This picture resembles microscopic hypersensitive response/apoptosis in an infected plant cell. The yellow arrow shows the location of the host nucleus. C) Polynuclearity was observed in a Pb-infected root at the elongation zone 16 days post-infection, compared to the mock healthy root control cells. This plant is tagged with GFP for nuclear envelope (ABRC stock number CS39988).

**Figure 3.**
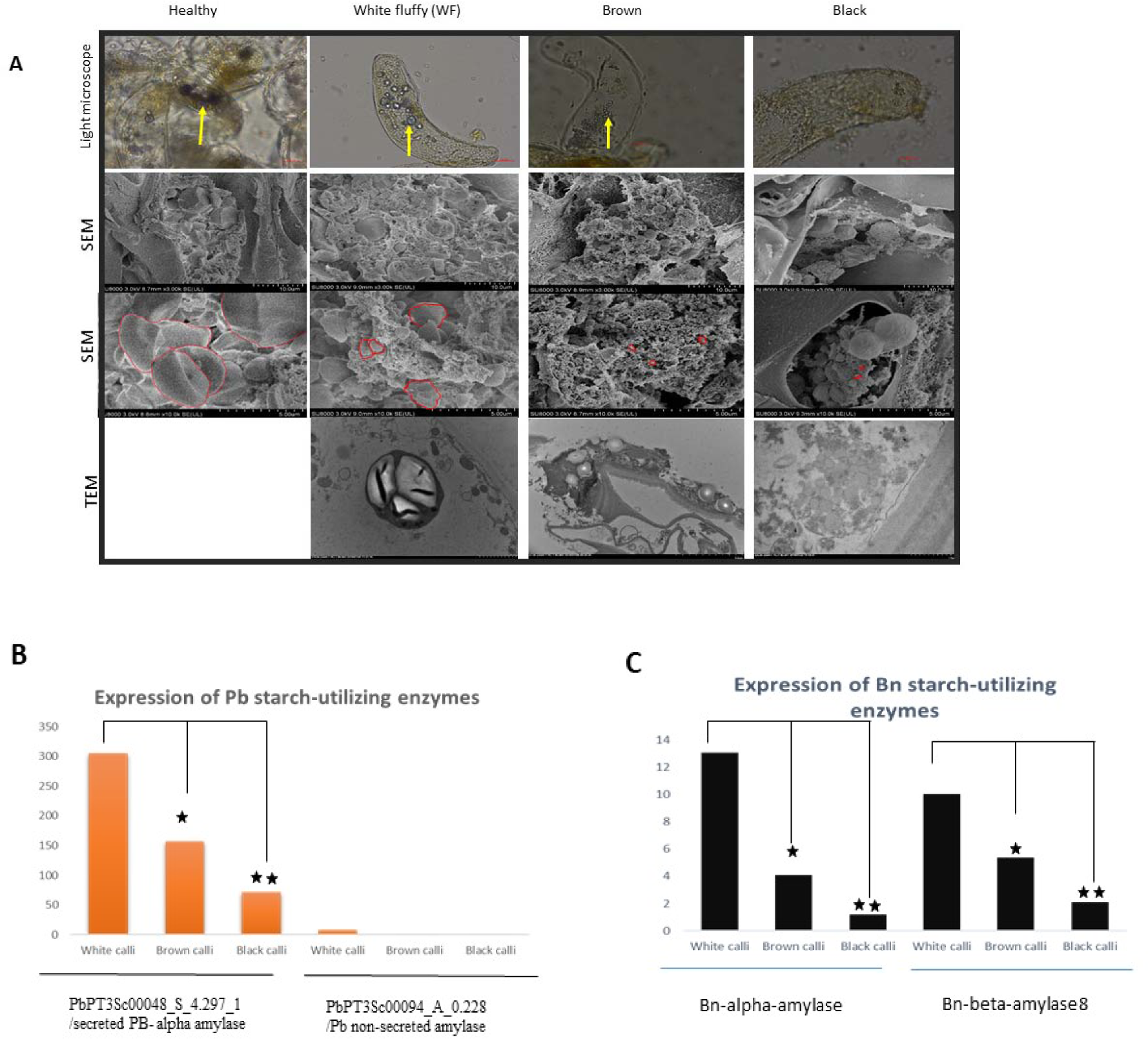
Starch accumulation and transcriptional activity of amylases in WF, Brown and Black calli of *Pb* axenic cultures as well as in different growth stages of *Pb*. A) Starch/amyloplast accumulation at different stages of *Pb* infection in axenic cultures. Top row from left to right is LM pictures of the starch granules accumulated in healthy, White fluffy (WF), Brown and black calli containing *Pb*, respectively. Starch granules were stained by lugol. Scale bar is 50 μm. The second row represents SEM micrographs of healthy, WF, Brown and Black calli, respectively. The scale bar is 10 μm. The third row is zoomed in view of the second row. Scale bar is 5 μm. The 4th row shows the TEM pictures of the amyloplasts in WF, Brown and Black axenic cell cultures. The starch granules are delineated by a red line around them. Transcriptional activities of starch-utilizing enzymes (amylases)are presented in B and C. B) Transcription of *Pb* alpha amylase is approximately 15 times higher than *Bn* alpha amylase, likely a secreted protein. *Pb*-amylase transcriptional activity is upregulated in WF cells, but it declines in brown and black calli possibly due to the shortage in cellular storage of the starch. PbPT3Sc00048_S_4.297_1 (*Pb* alpha amylase) is likely a secreted protein based on the presence of the N-terminal signal peptide, however, PbPT3Sc00094_A_0.228 lacks a signal peptide. The expression of the latter was slightly elevated in WF cells. C) Reduced expression of two candidate amylase genes of *B*. *napus* (*alpha amylase 1* and *beta amylase 8*) as measured by digital droplet PCR over time with the growth of calli (from young WF to older Black).

Relative expression of *B*. *napus* and *Pb amylase* genes correlated with the starch content of the cells and was highest in WF cells. Furthermore, our results indicate that starch was utilized by both the plant and the pathogen; although *Pb* seems to be the main user of starch as the expression of the secreted *Pb alpha amylase1* was over 30 times higher than *B*. *napus* amylases at all the stages of pathogen growth (Figure 3). As there were significantly bigger starch granules in WF cells compared to other cell types, the expression of *Pb* and *B*. *napus* amylase genes were also higher in WF cells indicating a higher need for starch in WF cells and suggesting that amylase activity is likely dependent on the abundance of its substrate. In other words, the breakdown of starch into smaller molecules of sugar is seemingly necessary for *Pb3* to exhibit a successful phase transition from its replication (zoospore and plasmodia formation) to feeding stages (enlargement of plasmodium cells).

Additionally, based on our observation, it is very likely that starch accumulation is induced by the pathogen, as we did not see the regeneration of the white calli from uninfected root cells, while the infected root calli cells regenerated fresh white cells that were filled with starch for pathogen consumption. These results are in agreement with the recent finding that suggests *Pb* is able to control sugar partitioning to induce a phloem sink in the local infected site (Walerowski et al. 2018).

### Oil biosynthesis and accumulation in *Pb* cells

It has been reported that oil droplets accumulate inside the *P*. *brassicae* zoospores (Aist and Williams 1971) and resting spores (Bi et al. 2016). Similar to the natural infection, we observed accumulation of oil bodies in the axenic cells. Staining of the cells with the lipidophilic fluorescent stain Nile red (excitation/emission maxima ∼552/636 nm) confirmed that there was little or no oil present in the healthy calli cells of canola roots; contrary to the accumulation of oil in the Pb-infected plant cells. Oil accumulation increased over time as indicated by the enhanced staining of the black cells compared to the brown and the WF cells (Figure 4). TEM of the infected calli, showed that oil bodies were only accumulated inside the parasite cells (Figure 5; Suppl Figures 4, and 5), including zoospore, young plasmodia, mature amoeba and resting spores. A considerable portion of oil bodies in the Pb cells consists of triacylglycerols (TGA) (Bi et al. 2016). Because the oil bodies were only found in the Pb cells, we questioned whether the TGA that was accumulated inside the Pb cells was synthesized solely by the Pb or the plant TGA biosynthesis machinery was also involved. Diglyceride acyltransferase, DGAT, catalyzes the formation of triglycerides from diacylglycerol in plants. The reaction catalyzed by DGAT is considered the last step in triglyceride biosynthesis, therefore expression of DGAT is a good marker for cellular TGA biosynthesis/accumulation. *B. napus* has four DGAT*1* genes designated *Bna*A*.DGAT1.a* (gene bank reference number JN224474), *BnaC.DGAT1.a* (JN224473), *BnaA.DGAT1.b* (JN224475), and *BnaC.DGAT1.b* (JN224476) that are involved in TGA biosynthesis (Greer et al. 2015). On the other hand, there are eight genes (with predicted signal peptide) involved in TGA biosynthesis/metabolism in Pb3 of which two were overexpressed at late pathogenesis i.e., 28 days post-infection (Rolfe et al. 2016; Suppl Table 3). The genes diacetylglycerol acyltransferase type 2a (*PbDGAT*; PbPT3Sc00016_Am_1.170), and phosphatidylglycerol phosphatidylinositol transfer protein (PbPT3Sc00086_Sm_1.204) were selected for expression analysis in *Pb3* axenic cultures. Elevated expression of *PbDGAT* and *BnDGAT* revealed that both the *B. napus-* and the *Pb* – *DGAT* genes are involved in TGA biosynthesis in clubroot infected cells (Figure 4). Furthermore, higher expression of *BnDGAT*, namely *BnaA.DGAT1.a* and *BnaC.DGAT1.a*, compared to *PbDGAT* genes suggested that triglyceride oil accumulation in pathogen cells primarily originates from the Plant. In agreement with Schwelm et al. (2016) who suggested that fatty acids are synthesized in plasmodial stage, expression of TGA marker genes was higher in the WF cells than the older stages calli grown from the infected cells (Figure 4), demonstrating that TGA biosynthesis is more prevalent in the early infection stage in both *Pb* and the host plant. Additionally, oil accumulation was clearly observed in two older stages of the axenic cultures (Figure 4; Suppl Figures 4, and 5), indicating that the oil bodies are likely synthesized, actively transported and accumulated in the pathogen cells.

**Figure 4.**
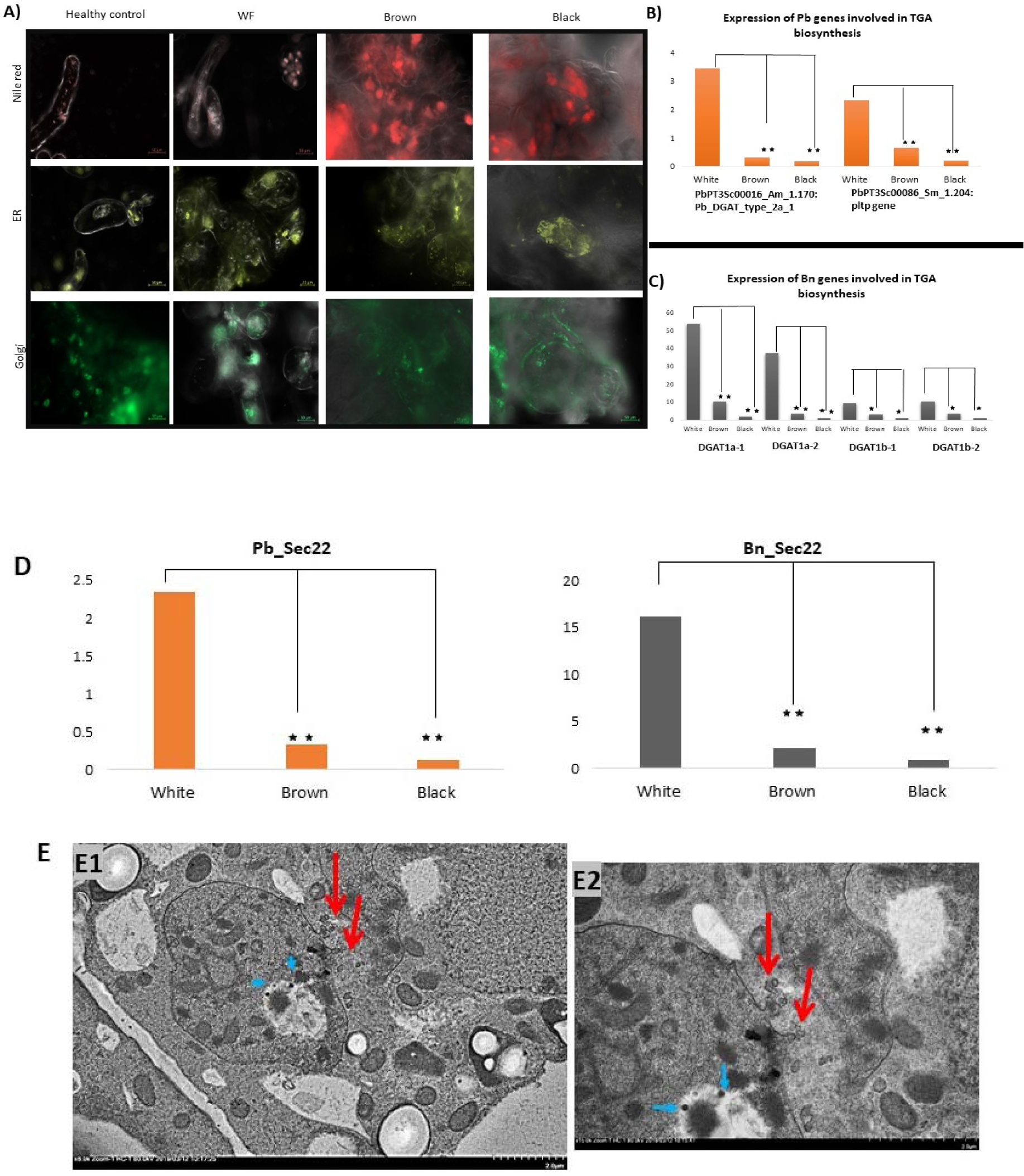
Accumulation of Lipids, Endoplasmic reticulum (ER) and Golgi inside *Pb* axenic cell cultures and transcriptional activation of candidate genes involved in Triglycerides biosynthesis as well as *Sec22* genes inside the cells of *Pb* and the host. A) Staining of the white fluffy (WF), Brown and Black axenic cell cultures compared to healthy control cells with dyes that are specific to Lipids (Nile red; top row), ER (Bodipy TR™ glibenclamide; middle row), and Golgi (BODIPY™ TR Ceramide complexed to BSA; third row). Pictures have been taken with dark field mode on a light microscope. Scale bars are 20 or 50 μm. There is hardly any lipid in healthy root calli cells –the residual red stain in these cells are due to the affinity of Nile red for lipid-bearing organelles. In general plant- and *Pb*-originated Lipid, ER and Golgi show substantial accumulation in the Black cells compared to other infected stages i.e., WF, Brown, and the healthy uninfected cells. Epifluorescent pictures were taken using Zeiss Apotome microscope. B and C show the expression of *DGAT* gene, a molecular marker for Triglycerides (TGA) biosynthesis in plants. B) Upregulation of the candidate *PbDGAT* genes in *Pb*-containing cells. *Pb_DGAT_type_2a_1*, as well as a *Pbpltp* genes are highly expressed in the WF cells suggesting *PbTGA* biosynthesis and intermembraneous transfer is mainly active in the earliest stages of the infection. C) Expression of DGAT genes of *Brassica napus* -infected with *Pb*. Higher expression of four *BnDGAT* genes in WF cells or early infection cells compared to other types of infected cells. Furthermore, higher expression of *BnDGAT* genes compared to *PbDGAT* gene strongly suggests that *BnTGA* biosynthesis is induced to a significantly higher level than that in *Pb*. D) Higher expression of both *PbSec22* and *BnSec22* suggests an active secretion activity inside the white cells. Both of these genes show less expression in the brown and black cells, compared to that in WF cells. Sec22 is active in vesicle trafficking, especially in the ER-Golgi protein trafficking. Higher expression of Bn_Sec22 compared to Pb_Sec22 suggests a major role for the Bn protein in ER- Golgi trafficking. E) TEM micrographs representing the involvement of the secretory system in Pb infection and in host defence. E1 shows the excretion of the *Pb* vesicles. *Pb* also engulfs and exerts lytic activities on plant defence proteins inside phagosome/phagolysosomes (blue arrows) in the plasmodia cell cytoplasm. A phagosome is a vesicle formed around a particle engulfed by a phagocyte via phagocytosis. E2 is an enlarged frame. Red arrows shows that the direct communication between *Pb* and host occurs in pseudopod-like papillae on the *Pb* surface. *Pb* excretes proteins through exocytosis into the host cytoplasm. In this figure, the electron density of *Pb* looks denser compared to the plant.

**Figure 5.**
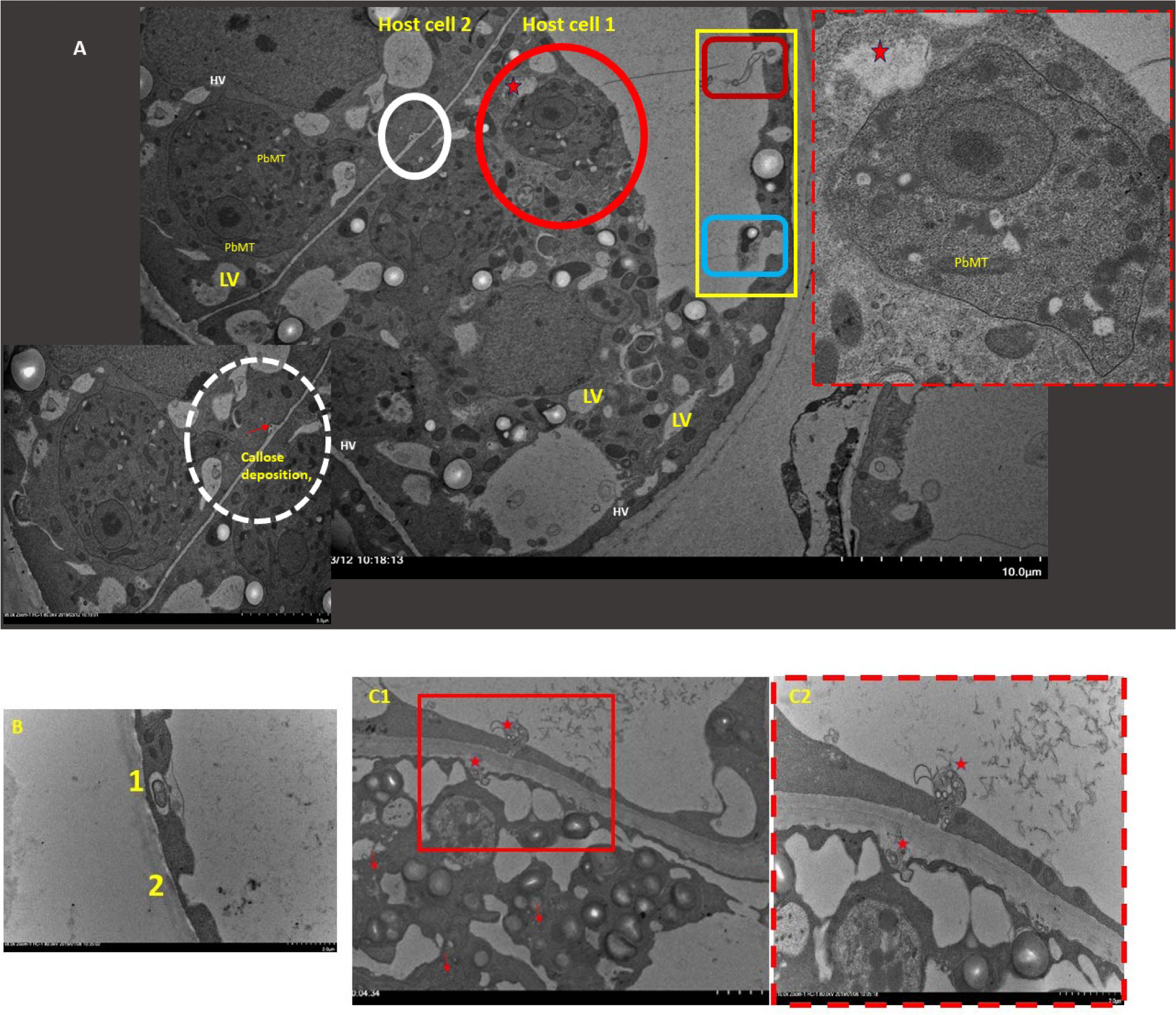
Plant defence mechanisms against Pb include autophagy, vacuolization, vesiculation, and callose deposition. Host cell activates its natural autophagy mechanism in defence response to infection by Pb. 5A is a TEM micrograph of two adjacent host cell containing multiple plasmodia or a combination of zoospores and plasmodia could be observed; scale bar is 10 μm. A) Upon penetration of the *Pb* zoospores, extensive vacuolation of the infected host cell is the secondary induced defence response to the infection. The yellow square shows how the plant cell utilizes an intravacuolar multivesicular compartmentation mechanism (Red rounded rectangle inside the yellow box) to sink the young zoospore inside the host lytic vacuole (Blue rounded rectangle). If a plasmodium encounters a plant autophagosome/ lytic vacuole (red asterisk) its cell wall disintegrates by the enzymes that are excreted outside the vacuole (red circle and its enlarged picture on the upright corner). The white circle (and the inset) shows the late papillae, and callose deposited on the host cell wall, where the plant plasmodesmata are prevalent. This is an indirect evidence of a *Pb* zoospore moving from the host cell 1 into cell 2. B) Biogenesis of a barrier around zoospore to trap the pathogen and sinking that toward lytic vacuoles is done in two stages i.e., 1. trapping/blockage and 2. Sinking into the lytic vacuole. C1) Multivesicular body (MVB) generation in a heavily infected cell and active excretion of its contents to the cytoplasm and the large vacuole of the neighbouring cell-This is a coordinated mechanism of autophagy and excretion by the plant vacuole. Active exocytosis, possibly Sec-22 dependent/coordinated exocytosis and endocytosis of MVB from infected cytoplasm to cytoplasm of an adjacent cell in WF cells. C2 is a magnification of the red square in C1. Red arrows are cross-section of host ER around Pb plasmodia. HV: Host vesicle; LV: Lytic vacuole.

### *P*. *brassicae*-induced adaptability of plant cell membranous organelles

There are few studies reporting morphological changes in *Pb*-infected plant cells but hardly any describe changes at the cellular level. Using axenic culture, we were able to observe alteration inside the plant cells as infection progressed. Notable changes were increased vesicle trafficking, induction of plant vacuole synthesis, increased size of plant cell nuclei and induced fragmentation of the nuclei or polynuclearity, as well as changes in the dynamic of the plant cell organelles including secretory system, mitochondria and plastids (Figures 2, 5, 6; Suppl Figure 3).

To support the growth of *Pb*, plant nuclei enlarge and show a drastic nucleo-cytoplasmic partitioning (Figures 2A, and 2C). Host nucleoli partitioning has been demonstrated to regulate cellular processes including hormone signalling, and plant-Microbe interactions (Allen and Strader 2021). The lack of immune cells in plants means each plant cell must act autonomously against an invading pathogen. All vegetative plant cells have vacuoles containing hydrolytic enzymes with a role in plant defence. We looked at vacuolar membrane dynamics upon *Pb* infection. In few events, the vacuolar membrane was collapsed as Pb infection progressed (Figure 5), similar to that in viral infection (Shimada et al. 2018.). Vacuolar membrane collapse is indicative of an effective plant defence response against viral infection resulting in a hypersensitive response (HR). Although the vacuole-dependent response to viral infection causes rapid HR, we observed gradual cell death in *Pb*-infected cells coinciding with the change in the thickness of infected cells (Figure 5) and their colour.

The secretory system in plant defence response involves multi-vesicular bodies (MVBs), lysosomes, exosomes, autophagy components, Endoplasmic reticulum (ER) and Golgi organelles. Pathogen-induced dynamics within plant endomembrane and membranous organelles have been observed in various interactions (Ruano and Scheuring 2020). We observed an overrepresentation of the secretory system in *Pb*-infected cells (Figure 4; Suppl Figure 4).

For instance, *Pb*-infected brown cells showed an accumulation of MVBs (Figure 5). On the other hand, pathogens have evolved effectors to prevent regular trafficking within infected cells through subverted autophagy and MVBs, believed to be the plant’s natural mechanisms for effective defence activation (Ruano and Scheuring 2020). Attachment of MVB with the plasma membrane and ER is part of unconventional trafficking inside the infected cells to counteract pathogen attack. Additionally, lysosomal exocytosis leads to the secretion of lysosomal content upon lysosome fusion with the plasma membrane. MVBs deliver their contents extracellular or into lysosomes.

Another vacuole-dependent defence response which was observed by EM is autophagy (Figure 5). Autophagy is represented as a natural expression and completion of a membrane around pathogen-induced particles within the cytoplasm. Autophagy causes the production of tiny vacuoles. This kind of vacuole is known as autophagosome which after merging into lysosomes results into the breakdown of the infection-induced particles. Autophagy degrades and removes unnecessary components through a lysosome-dependent mechanism. Empty lysosomes are observed as small vacuoles (Figures 5 and 6). Induced autophagy in *Pb*-infected cells is always accompanied by significant compartmentation of the plant cell cytoplasm. Cleavage in the infected cell cytoplasm could by itself be assumed as an effective defence mechanism against Pb which resides within the cytoplasm to feed and multiply.

**Figure 6.**
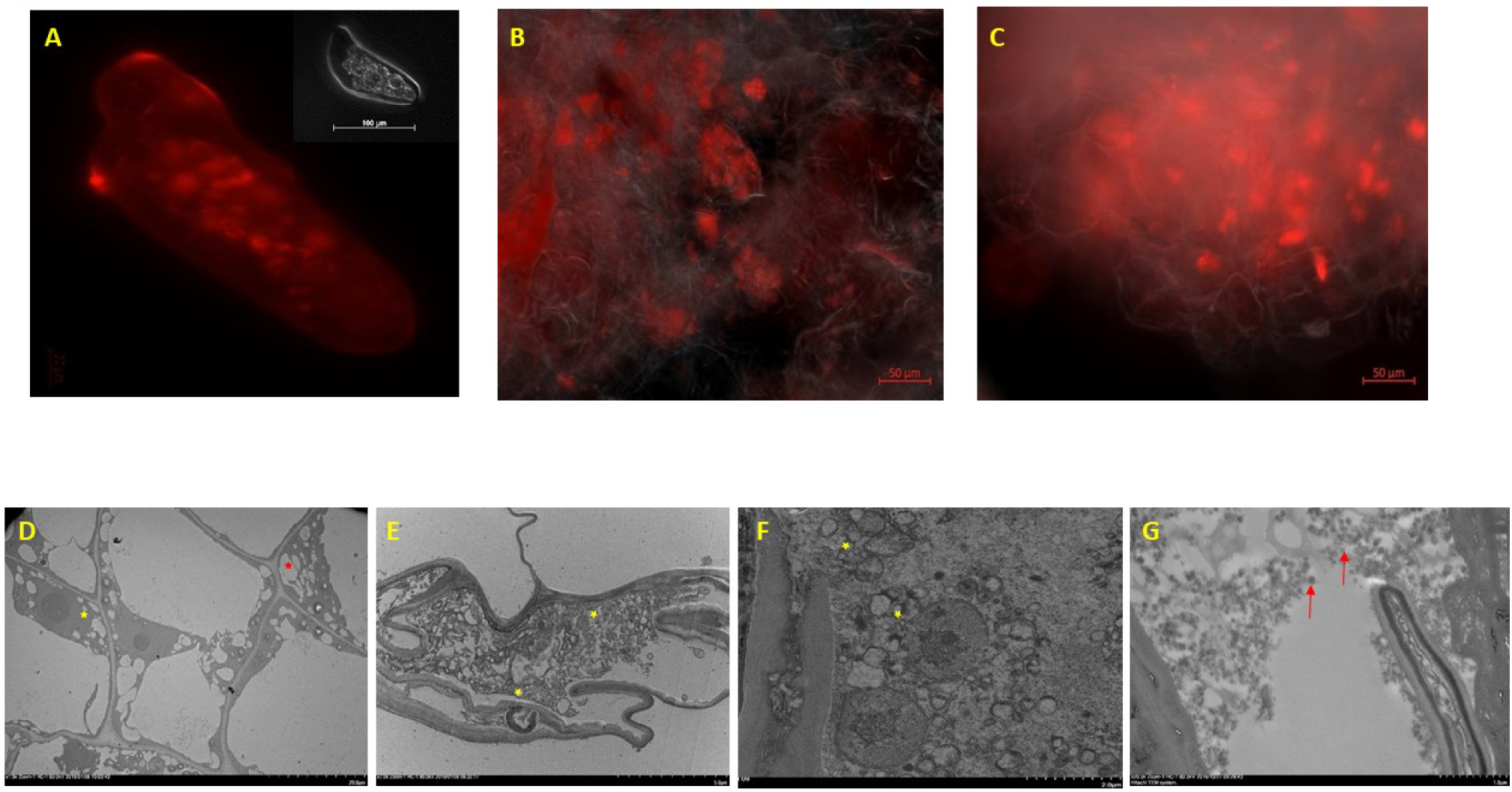
Extreme vesiculation of *Pb*-infected cells. Vesiculation of single WF cell (A), brown (B), and black cells (C) stained with fluorescent dye FM4-64 and observed with epi fluorescens light. In the upright corner of picture A, a single WF cell is shown under the darkfield LM. Figures D to F are TEM micrographs of the Pb-infected WF (D), Brown (E), and Black (F) cells. D) Plant WF cells that were just infected were induced for accelerated lytic vacuole (red asterisk) generation as well as vesicles (yellow asterisk). An increase in the of plant vacuole quantities help the cell authophagy in Pb-infected cells. G) Clathrin-covered vesicles in Black Pb axenic cells. Clathrin-coated vesicles (CCVs) are observed in the Black cells (red arrows). CCVs mediate the vesicular transport of cellular cargo via the trans-Golgi network, connecting endosomes, lysosomes, and the cell membrane. FM4-64 stains vesicles and over time it spreads throughout the vesicular network from the plasma membrane to the vacuole, including the components of the secretory pathways. Vesicles (yellow asterisk).

As a plant defence mechanism, *Pb* cells are contained in plant-originated vacuoles in either zoospore or plasmodium stages in the infected cells. After penetration of the *Pb* zoospores into the infected cells, autophagy seems to be a secondary induced response to the infection following the extensive vesiculation of the host cell. We also observed that when a plasmodium comes in contact with an autophagosome/ lytic vacuole, its cell wall disintegrates by the lytic vacuole (Figure 5). To survive the mature plasmodium has to avoid the plant vacuoles while growing inside the host cytoplasm. In addition, the infected plant cell utilizes an intravacuolar multivesicular compartmentation mechanism to detach and sink the young zoospores inside the host main lytic vacuole (Figure 5). An increase in the quantities of plant vacuole helps the autophagy in *Pb*-infected cells. Lastly, the generation of Clathrin-coated vesicles (CCVs) in the black cells (Figure 6) mediates the vesicular transport of cargo such as proteins between organelles in the post-Golgi network connecting the trans-Golgi network, endosomes, lysosomes, and the cell membrane.

The endoplasmic reticulum (ER) and the Golgi apparatus are extremely dynamic organelles which have critical roles in various cellular processes (Chatre et al. 2005). Live imaging of the axenic cell cultures stained with fluorescent Bodipy™ stains that selectively bind to ER (ER-Tracker Red (Bodipy TR™ glibenclamide, Invitrogen), or Golgi (BODIPY™ TR Ceramide complexed to BSA, Invitrogen) revealed that ER and Golgi apparatus are accumulated in the *Pb*3-infected cells (Figure 4). This was supported by our TEM studies showing accumulation of ER around the plasmodia (amoeba cells) in axenic culture (Figure 5). Additionally, we observed a clear accumulation of smooth and rough ER as well as Golgi bodies inside mature plasmodium (Suppl. Figure 4). Cell membranes of the plasmodium were thinner in some spots, where the *Pb* and plant ER seem to communicate. The presence of the hub of direct interaction could be an indication that clubroot pathogen makes use of the plant secretory system to secret its proteins into the host cell.

In eukaryotic cells including plants and animals the molecular machinery involved in the secretory pathway appears to be highly conserved. Soluble N-ethylmaleimide-sensitive factor attachment protein receptors (SNARE) proteins including Sec22 are involved in cargo vesicle trafficking and induction of membrane fusion (Yang et al. 2018). To understand whether the ER accumulation was accompanied by transcriptional activation of the genes involved in ER morphology, we studied the expression of *Sec22* genes of the host (*B*. *napus*) and the pathogen (Pb). *AtSec22* has been shown to be involved in protein trafficking at the ER-Golgi interface (Chatre et al. 2005). Sec22 also regulates ER morphology by regulating ER-Golgi trafficking (Zhao et al. 2015). Higher expression of both *PbSec22* and *BnSec22* suggested intensified active secretion inside the cells (Figure 4). It has been shown that other fungi such as *Colletotrichum orbiculare* secrete their virulence factor to a Biotrophic Interface (BIC) at the primary hyphal neck via SeC22-mediated trafficking coupled with exocytosis (Irieda et al. 2014). Expression of *BnSec22* and *PbSec22* was significantly higher in WF cells and gradually declined in the infected cells as infection progressed as the infected calli turned brown and black in colour (Figure 4). In agreement with our TEM micrographs where we observed an active excretion in the newly infected cells, Sec22-mediated traffic could be accounted for the significant exocytosis activity in WF cells. This process could help the cell to excrete the pathogen-produced effectors and lytic enzymes to be transported into neighbouring cells (Figure 4).

Additionally, the dynamic of plant cellular components in the early stages of infection was studied using a transgenic Kaleidocell Arabidopsis plant (Kato et al. 2008) which was tagged with several spectrally different fluorescent proteins. Nuclei, plastids, mitochondria, and plasma membranes were tagged with cyan, red, yellow, and green fluorescent proteins, respectively. Nuclei were not clearly observed in the roots of Kaleidocell plants; however, our results showed a strong accumulation of plant mitochondria as well as constraints in the plasma membrane at the site of zoospore penetration –mainly cell corners- in the epidermal cells (Figure 2), suggesting that mitochondria accumulation is one of the earliest responses of the plant cells to infection by clubroot. Our TEM studies of the axenic cultures in newly infected WF cells also clearly showed accumulation of mitochondria around the zoospores inside the cytoplasm of the plant cell (Figure 2). Additionally, an abundance of plant cell mitochondria was always observed around growing amoebae cells (Figure 5). The Amoebae cells contain their own mitochondria as well (Figure 5) which is likely to meet the need of the growing pathogen for energy. Moreover, plastids accumulated in the infected as well as in the neighbouring cells. Accumulation of actin filaments around the infection site was also observed (Figure 2), which suggests the restructuring of actin in growing cells.

### Active intercellular movement of *Pb*: association with Plasmodesmata

Microscopy observation revealed that zoospores were present in WF cells and they freely swarm inside the cytoplasm (Suppl Figures 5 and 6). We investigated the possibility of whether the zoospores that are generated on agar plates inside axenic cell cultures could potentially infect neighbouring. Live cell microscopy showed constraints in the cell wall and cytoplasm of the Arabidopsis root cells infected with *Pb*. At the same time, we studied callose deposition in the cv. Granaat root cells infected with zoospores that were released inside the RS-containing black calli. TEM analysis revealed extensive callose deposition on the cell wall of the WF cells (Figure 5; Suppl Fig. 6). Callose deposition was also visualised by Aniline blue and Calcofluor staining at days 3-7 post-infection by *Pb* on the cell walls adjacent to other plant cells as well as to the peripheral sides of the cortical cell layer, suggesting that the secondary zoospores are likely to pass through the cell wall to the neighbouring cells. Callose deposition on all the cells suggests that plant uses this mechanism to block the movement of zoospores to the adjacent cells (Suppl Figure 6). Callose is a glucose-based polysaccharide with essential roles in various structural changes in plant cells (Slewinski et al. 2012). Localised callose deposition known as papillae is a typical defence response to pathogen attack. Our TEM experiments revealed that there were callose depositions as well as papilla formation on the cell walls of the WF calli, suggesting a plausible movement of zoospores through the cell walls of the *Pb3* infected axenic cultures (Suppl Figure 6, and Figure 5A).

Plasmodesmata regulate the symplastic intercellular trafficking and act as well-regulated openings for cytoplasmic trading of pathogens such as viruses. Plasmodesmatal trafficking could be either passive or active based on environmental stimuli (De Storme and Geelen 2014). Callose turn-over regulates the passage through the plasmodesmata gates. Figures 2A shows the active movement of a young plasmodium between a *Pb*-infected cell to its adjacent cell through transmembrane.

The tight association of the *Pb* zoospores and plasmodia with plasmodesmata openings suggests that zoospores and plasmodia are possibly able to traffic between the cytoplasm of the neighbouring plant cells. The movement is likely to involve hijacking the plant’s cellular exocytosis through plasmodesmata. Six different protist species were shown to traverse through narrow Channels (Wang et al. 2005). All the species tested were able to pass through channels at different speeds based on their movement strategy and the size of the pores.

Additionally, *Pb* seems to multiply during the plant cell mitosis as observed in stage 1 cells (Suppl Figure 7). Our results indicates that the zoosporangia inside WF cells could be incorporated into the newly formed cells during plant cell mitosis thus carrying the pathogen over to the new cells. TEM micrograph strongly suggests that other roots of trafficking in plant cells are hijacked by *Pb* to move from cell to cell. Suppl Figure 7 shows presence of porosomes in the pb infected cells pointing to the role of these structures in cell-to-cell movement of Pb. Porosomes are cup-shaped structure and docking location for the secretory vesicles. These means of intercellular trafficking have not been reported previously for *Pb*. Furthermore, the movement of *Pb* is favoured by both induced and passive intercellular mechanisms including but not limited to induced cell mitosis and hyperplasia, and porosomes. Prosomes are funnel shape appendages on the cell surface that open up into the natural openings of the cell (Suppl Figure 7).

## Discussion

*Pb* as a unicellular organism, must meet all the biological functions necessary for its virulence and completion of its life cycle that is achieved by many different specialized cell types in multicellular organisms. In order to cope with the harsh plant intracellular environment, *Pb* changes its cellular-spores-morphology over the course of infection. *Pb* is known to alter plant Auxin and Cytokinin homeostasis (Rolf et al. 2016). We observe increase in in cell number and cell size in infected calli, possibly due to changes in the balance between cytokinin and auxin induction.

Mechanism of *Pb* penetration and entry into the root cells has been studied by Aist and Williams (1971). However, there has not been in-depth study on cellular changes of *Pb* and its host plant during infection. Most of our insights into *Pb*-Plant interactions at the cellular level were aided by the in-vitro axenic cell culture method optimised for this study. Addressing how dynamics of the plant cell organelles contribute to cellular homeostasis in response to *Pb* infection is exceptionally challenging. We studied the expression of a few candidate *Pb* effectors including *PbBSMT*, *PbDGAT*, and *Pb amylase* in different stages of the axenic cultures as a few examples to show that *Pb* effectors are transcribed under the axenic culture growth similar to natural root infection.

By applying live imaging and electron microscopy, we found similarities between *Pb* and the model human parasite *Plasmodium falciparum* infectious structures inside the infected cells. As an example, the life cycle of *Pb* which consists of distinct steps including “invasion-replication-feeding-replication-egress” resembles the infection process reported for *P. falciparum* in human erythrocyte (Tilly et al. 2011). *P. falciparum*, the causative agent of malaria, completely remodels the infected human to acquire nutrients and to evade the immune system. To do this, the parasite secrets more than 10% of all its proteins into the host cell cytosol. There is little known about the plant changes at the cellular level after infection with *Pb*.

Our results suggest the induction of drastic and orchestrated changes in *Pb*-infected cells varying from starch metabolism, oil accumulation, and active alteration in the kinetics of plant intracellular organelles. Interestingly, we found that although oil bodies only accumulate inside the *Pb*-infected root cells, both plant and parasite *DGAT* genes are highly expressed in the young infected calli (white fluffy cells), which indicates a need for the oil accumulation at this stage which is a switch between replication and growth stages of the pathogen. Transcriptional studies of *Pb* infection by Schwelm et al. (2016) suggested that fatty acids are synthesized in the plasmodia and degraded in the resting spore. The higher expression of *B. napus DGAT* genes in the infected axenic cells indicates that in addition to the pathogen’s own oil biosynthesis, *Pb* relies on the plant as the major resource for oil accumulation. It has been suggested that lipid biosynthesis could act as a membrane shield protecting *Pb* from the host defence while coexisting inside the host cell, in addition to its role in the suppression of plant immunity against fungal pathogens (Basso et al. 2022).

Lipids are not readily transferable throughout the cell membranes, therefore the expression of PbPtlp protein seems to catalyze the intermembrane transfer of phosphatidylglycerol and phosphatidylinositol, by which TGA transfer would be facilitated between the membranes of *Pb* and *Bn*. These results demonstrate a need for plant-based biosynthesis of TGA, and pathogen-based catalysis and the transfer of TGA in *Pb*-infected cells.

Oil accumulation seems to be a key factor in the hypertrophy of plant root cells. Adding 10% glycerol to the roots of Arabidopsis plants on Murashig and Skoog (MS) media resulted in a faster appearance of galls on the roots. Although the roots were thicker in infected Arabidopsis than the roots of infected Arabidopsis without glycerol (control-not shown).

Utilization of starch granules by *Pb* cells occurred at a higher rate compared to infected plant cells. These changes correlated with the expression of a secreted *Pb amylase* gene which was significantly higher (over 75 to 100 times) than the plant *amylases*. Walerowski et al. (2018) showed that Pb forms a feeding relationship with its host through the induction of SWEET Sucrose Transporters within developing galls. Our results demonstrate that sugar metabolism is an active mechanism that is essential for *Pb* growth and replication.

*Pb* is an intracellular pathogen and needs to be protected from the host harsh intercellular environment triggered by plant immunity response. We show that *Pb* infection induced the accumulation of vesicles inside the cells. Increased budding (vacuolization), induced exocytosis and autophagy in *Pb*-infected cells seem to be examples of the host cellular defence. Autophagy for example acts as a cytoplasmic delivery of pathogen- and plant-derived components to lysosomes for digestion. Budding, on the other hand, is membrane-originated and involves enzymatic digestion of proteins such as pathogen effector proteins. To alarm neighbouring cells, exocytosis acts as a cell-cell communication tool. Vacuolization after internalization of the intracellular human pathogens is a mechanism which is used by some bacteria (Sun et al. 2018). Intracellular bacteria would benefit from endocytosis/vacuolization as they become protected from exposure to defence mechanisms inside the cell. This process would end by releasing the pathogen-containing vacuoles through exocytosis. *Pb* internalization may follow a similar mechanism.

Another major defence response against pathogens including *P*. *brassicae* is callose deposition at the early stages of infection and thickening of the cell wall at the later stages. *Pb* would change its lifestyle from being parasitic with little or no nutrient requirements- to feeding stage i.e., plasmodium cells and finally forms resting spores. In other words, *Pb* development in infected roots occurs in two steps i.e., replication and growth. Primary zoospores would lose their flagella soon after their contact with root surface. However, the secondary zoospores maintain their flagella while inside the cell.

Plant defence response to Pb comprises various mechanisms including, but not limited to production of lytic vacuoles, and suberification of cell wall to inhibit the movement of the pathogen to the neighbouring cells through dynamic callose deposition.

Dynamic regulation of plant endomembrane system in response to PTI has become the focus of host-pathogen interaction in recent studies. The plant defence against *Pb* consists of preformed defence barriers and plant immune response including secretion of antimicrobial proteins. One of the processes used by the pathogen to counteract the host defence is to manipulate vesicle trafficking. However, plants have evolved another layer of defence in response to plant immunity which includes the formation of ER bodies or fusion of vacuoles with PM (Ruano and Scheuring 2020). The intercellular movement of some plant pathogens including viruses (Lee, 2014), as well as the multicellular fungal pathogen, *Magnaporthe grisea*, (Kankanala et al. 2007) has been well documented. *M*. *grisea* appears to kill any invaded plant cell as it moves into the adjacent cells, however, *Pb* seems to move with the least harm to the infected cells.

Transmission electron microscopy showed *Pb* zoospores are favourably accumulating next to plasmodesmata-where natural pit fields (sizes varying between 50 nm to 5 um wide)- are present between neighbouring cells. *M*. *grisea* could readily pass through the natural holes in the plant cells that are up to 300 nm wide. Interestingly, Wang et al. (2013) showed that, similar to plant response to *Pb* which is mainly Salicylic acid (SA)-dependent, SA signalling plays a critical role in the regulation of cellular networks via plasmodesmata as part of their defence response. Additionally, callose deposition exerts physical constraint on plasmodesmata, hence restricting the traffic through the channels. These evidence strongly suggest that *Pb* co-opts plasmodesmata for cell-to-cell movement. This structure resembles intercellular movement of multivesicular bodies (MVBs) at the penetration site of powdery mildew-barley pathosystem. However, MVBs do not induce callose and they normally originate from lytic vacuoles (An et al. 2006).

In order to expand biotrophic invasion into the neighbouring cells, other strategies than direct entry through force by the means of satchel are employed by *Pb*. As an example, our work revealed other novel modes of intercellular movement of *Pb*, i.e., induction of cell mitosis, and movement through porosomes.

This paper for the first time describes in-depth changes at the host cellular level in response to *Pb*. *Pb* changes the cell size leading to the biogenesis of root galls. In order to infect the plant cells, *Pb* actively alters the size and the integrity of the nucleus, and plant vacuolar and secretory systems. To counteract plant uses its natural secretory system to combat the pathogen. Our study also reveals novel ways that *Pb* employs to colonize the root cells through moving from one cell to the neighbouring cells.

## Material and Methods

### Axenic culture of Pb

Axenic cultures were prepared according to Bulman et al. (2011). Briefly, the pure resting spores from various clubroot pathotypes including Pb2, 3, 5, 6 and 8 that were received from Dr. S.E. Strelkov, University of Alberta, were used to inoculate the roots of *Brassica napus* cv. DH12075 and *Brassica rapa* ssp. *pekinensis* cv. Granaat. After 6 to eight weeks the roots with the symptoms of clubroot were harvested and disinfected. The infected roots as well as the healthy control roots were cut into small pieces and incubated at 24 C in dark on MS Agar plates. Six weeks later the calli, regenerated from infected roots, were selected and stored on new agar plates. Every six weeks calli were transferred into new agar plates.

Infectivity of the axenic cell cultures was examined by inoculating cv. DH12075, cv. Granaat and Arabidopsis thaliana Col-0 accession as described by Djavaheri et al. (2018) using 2 × 10^7^ ml^−1^ resting spore (RS). Gall formation was evaluated at six weeks post-inoculation.

### DNA extraction and sequencing

Total DNA was extracted from infected calli using CTAB, and then sequenced using illumina MiSeq (Illumina, CA, USA). All the sequence reads were mapped to the reference genome (Rolfe et al. 2016) using CLC (Version 8.1.1, CLC Bio, Aarhus, Denmark).

### Plant materials

Plants were grown on a soilless potting mix and were kept in a growth cabinet with 8 h of light (100 μmol/m2/s) at 22°C (light) and 20°C (dark) and 70% relative humidity for Arabidopsis and 18 hrs light for brassica plants. Transgenic Kaleidocell Arabidopsis plants as well as the Arabidopsis Col-0 harbouring GFP-WIT1 for nuclear envelope (ABRC stock number CS39988) have been described elsewhere (Kato et al. 2008; Zhao et al. 2008).

### Staining the calli and microscopy

Live cell imaging of the Axenic cultures was done using Zeiss (Oberkochen, Germany) light or epifluorescent microscopes. The images were processed using ZenBlue® software according to the manufacturer. Epifluorescent microscopy was conducted using FM™ 4-64 Dye (N-(3-Triethylammoniumpropyl)-4-(6-(4-(Diethylamino) Phenyl) Hexatrienyl) Pyridinium Dibromide) that selectively binds to vacuolar membranes with red fluorescence (excitation/emission maxima ∼515/640 nm). This lipophilic dye is an important tool for visualizing vacuolar organelle morphology and dynamics, for studying the endocytic pathway. Staining of the cells with the lipidophilic fluorescent stain Nile red (excitation/emission maxima ∼552/636 nm), was used to study the accumulation of oil bodies in the infected plant cells (Bi et al. 2016).

Additionally, Lugol and Calcofluor -(excitation/emission: ∼365/ 397 nm)- (Herburger, and Holzinger, 2016) were applied to stain the starch granules and callose deposition in the live cells. Furthermore, live imaging of the axenic cell cultures stained with fluorescent bodipy™ stains that selectively bind to ER (ER-Tracker Red-Bodipy TR™ glibenclamide, Invitrogen), or Golgi (BODIPY™ TR Ceramide complexed to BSA, Invitrogen) was used to study the accumulation of ER and Golgi apparati in the infected cells.

Transmission electron microscopy (TEM) was performed as follows. Specimens were post-fixed in Osmium tetroxide (1%) in 0.1M Sodium Cacodylate buffer for 1 hr at room temperature followed by 2 rinses in distilled water to remove traces of Osmium. Specimens were dehydrated through graded ethanol series (50%, 70%, 85%, 95%, and three times 100%), then embedded in LR white resin and polymerized at 60 degrees for 24 hours. Specimens were subsequently sectioned on a Diatome diamond knife at a thickness of 90nm and mounted on a 200-mesh copper grid. Specimens on grids were post-stained for 20 mins with 2% Urnyl acetate, rinsed then stained for 10 minutes with Reynolds Lead Citrate. Sections were observed using a Hitachi HT7700 Digital Transmission Electron Microscope.

For scanning electron microscopy (SEM) various cell types were fixed in 2% glutaraldehyde (GA) in 0.1M Sodium Cacodylate (NaCAC), pH 7.2. After fixing for a few hours at room temperature, samples were moved to a fridge. On the following day, specimens were washed with 0.1M NaCAC and subsequently stored in the fridge.

Then dehydration was performed in 30,70, and 70% ethanol for 20 min. each. Following overnight incubation in the fridge white fluffy (WF) as well as brown and black calli were sectioned with a razorblade on the following day, and then dehydration continued in 80, 90, 95 and three times 100% ethanol. Subsequently, the critical point drying run (Polaron E3000) was performed. Samples were then coated with 10nm gold (Quorum Q150T ES). The Scanning electron microscopy was done using SU8010, Hitachi High Technologies (Tokyo, Japan).

### Gene expression analyses

RNA was extracted using Purelink RNA mini kit (Ambion, Invitrogen, MA, USA) from four samples that were taken from each stage of the infected calli. cDNA synthesis was done according to the manufacturer (iScript RT Supermix, BioRad, CA, USA). To quantify the expression levels of various genes, droplet digital PCR-ddPCR- (Whale et al. 2012) was employed as described (Djavaheri et al. 2019). Expression of the genes responsible for TGA biosynthesis, starch metabolism (alpha and beta-amylase), *PbBSMT* (Djavaheri et al. 2019) as well as *Sec22* in *B*. *napus* and *P*. *brassicae* were compared to the expression of *B. napus actin 2* (*BnActin*; Bahar et al. 2018). A list of the genes, primers and probes for ddPCR is presented in Supplementary Table 2.

Relative expression of the genes responsible for TGA biosynthesis in *B*. *napus* including *BnDGAT1a-1*, *BnDGAT1a-2*, *BnDGAT1b-1* and *BnDGAT1b-2* (Greer et al. 2015) were studied in WF, Brown and black calli. Additionally, the expression of selected Pb genes including *PbBSMT* (Djavaheri et al. 2019), *Pbalpha amylase 1* (PbPT3Sc00048_S_4.297_1, hypothetical secreted protein), a non-secreted Pb amylase likely involved in hydrolyzing gelatinized starch (PbPT3Sc00094_A_0.228), *PbDGAT2a* (the predicted secreted protein diacylglycerol acyltransferase type2a; PbPT3Sc00016_Am_1.170), a Pb phosphatidylglycerol phosphatidylinositol transfer protein (Ptlp: a non-secreted phosphatidylglycerol phosphatidylinositol transfer protein PbPT3Sc00086_Sm_1.204) and (*PbSec22*), were studied relative to *BnActin*. From the list of the Brassica genes that were discovered to be involved in lipid biosynthesis (Bi et al. 2016), the expression of *PbDGAT2a* and *Pltp* were increased over time in Pb3-infected roots (Rolfe et al. 2016).

## Acknowledgement

Authors thank the following individuals for technical help: C. Coutu, Agriculture and Agri-Food Canada; Sobchishin, Larhonda, and Eiko Kawamura, the Western College of Veterinary Medicine, University of Saskatchewan for their technical assistance in conducting TEM, and SEM experiments. Authors would like to thank Dr Mark Smith, Agriculture and Agri-Food Canada- for helpful discussion on the oil biosynthesis in *Brassica* plants.

