## supplementary file for "Plasticity of host cell organelles in response to Pb infection"

### Supplementary Materials:

Suppl. Table 1. Illumina Miseq confirmed the presence of the pathogen within the plant cell with over 10% of the total sequence reads being from *Pb3*.

| | Count | Percentage of reads |
|---|---|---|
| Mapped reads to Pb3 | 1,820,145 | 11.4% |
| Un-mapped reads | 14,251,799 | 88.68% |
| Paired reads | 1,741,494 | 10.84% |
| Broken paired reads | 78,651 | 0.49% |

Suppl. Table 2. Primers and probes used in droplet digital PCR experiments to study expression levels of various genes of *Brassica napus* or *Plasmodiophora brassicae*.

| Gene | Gene name/function | forward primer | reverse primer | probe |
|---|---|---|---|---|
| PbPT3Sc00026_A_1.308 | PbBSMT/Secreted | CCCGCATGTTCTATGTCATTCC | CGGTTAGCGTTCGTTCCATAAA | ACATCTTGTCGACGTATTGCCCGT |
| PbPT3Sc00048_S_4.297 | Pb-alpha amylase/hypothetical secreted | CCCAATCCGCAACATGAGTA | GTGTACTAGGAGAACGAGAAACG | TCGCTGTATACATCCGGGCACAAC |
| PbPT3Sc00094_A_0.228 | Pb amylase-non-secreted/non-secreted amylase | CGCACGTACTGGAACGATTT | ATGTCGTTAACGCGGCTATG | CGATCACGCTGAAGATGTGCTCCT |
| PbPT3Sc00016_Am_1.170 | PbDGAT2/hypothetical secreted | GCGTCTTCGTTTCGGTTCT | TCGGTTGTGTCGATGGTTTG | CTTCTCTGCGTCTCGTCATCGGT |
| PbPT3Sc00086_Sm_1.204 | Pltp | GGTCTCAAAGGCCATCGATATT | CGCATGTGGTACTCGAACT | CCGCAGTCCACGGTTGTATTCTGT |
| PbPT3Sc00039_A_6.347_1 | PbSec22/non-secreted | CAACATCGCCAAGCTGAATG | CCGTTTGAGGATCTCGTTCATA | ACACAGGTGCACAATGTTATGAGGTCA |
| JN224474.1 | BnDGAT1a-1 | GATTCTGGAGGCGTCACTATG | AAGAAGTCCGTTGGAGGAATC | TCGATACGCTTCGTCACCGGAAAC |
| JN224473.1 | BnDGAT1a-2 | GATTCTGGAGGCGTCACTATG | CAGGAAGAAGTCCATTGGAAGA | TCGATACGCTTCGTCACCGGAAAC |
| JN224475.1 | BnDGAT1b-1 | CGAAAGGGATCGGGTTGATT | GATTCCCTAGTTTCGGCATCTC | TAAATTCGCTGTTCCCTGAGCCTCC |
| JN224476.1 | BnDGAT1b-2 | CTCAGGGAACAGCGAACTTAG | GGCCGATACGTAAACCTTACA | TGGCGGAGATACCGAAACTAGGGA |
| NC_027757.2 | Bn alpha amylase 1 | GACCTTAGCAACTCTGGAATCA | TGAAAGTCAAGGAACGGGATAG | TGTCTGGCTTCCTCCTCCTTCTCA |
| NC_027768.2 | Bn beta amylase 8 | GAGAGAGGGAGAAGGAGAAAGA | GGAAGTTACCGTACTGTCTCAAC | AAAGACACAGGCGAGCCATCACTA |
| NC_027762.2 | BnSec-22 | TTGCATATCTCCCTGGATCAAC | ACCTGATACGAGTGTTCCAATG | ACCCGGTTCTCAGATCCTGAGAAGA |

Pltp: a Pb phosphatidylglycerol phosphatidylinositol transfer protein ; ns:non-secreted, hs:hypothetically secreted

Suppl.Table 3. List of secreted *PbDGAT* genes of *Pb3* and their expression on days 21 and 28 dpi on Arabidopsis infected roots.

| | days post Pb3 infection | | |
|---|---|---|---|
| **Pb gene** | **21 dpi** | **28 dpi** | **Function** |
| PbPT3Sc00003_A_0.276 | 5.290 | 4.497 | acyl-sn-glycerol-3-phosphate_acyltransferase_pls1_1 |
| PbPT3Sc00016_Am_1.17 | 1.312 | 3.884 | diacylglycerol_acyltransferase_type_2a_1 |
| PbPT3Sc00018_Sm_3.255 | 5.323 | 6.722 | triacylglycerol_lipase_1 |
| PbPT3Sc00057_Am_1.89 | 4.057 | 2.654 | glycerol-3-phosphate_dehydrogenase_1 |
| PbPT3Sc00060_Am_0.151 | 2.116 | 3.830 | diacylglycerol_acyltransferase_family_protein_1 |
| PbPT3Sc00086_Sm_1.204 | 5.074 | 7.039 | phosphatidylglycerol_phosphatidylinositol_transfer_protein_1 |
| PbPT3Sc00092_A_3.279 | 2.259 | 4.004 | triacylglycerol_lipase-like_protein_triacylglycerol_lipase_1 |

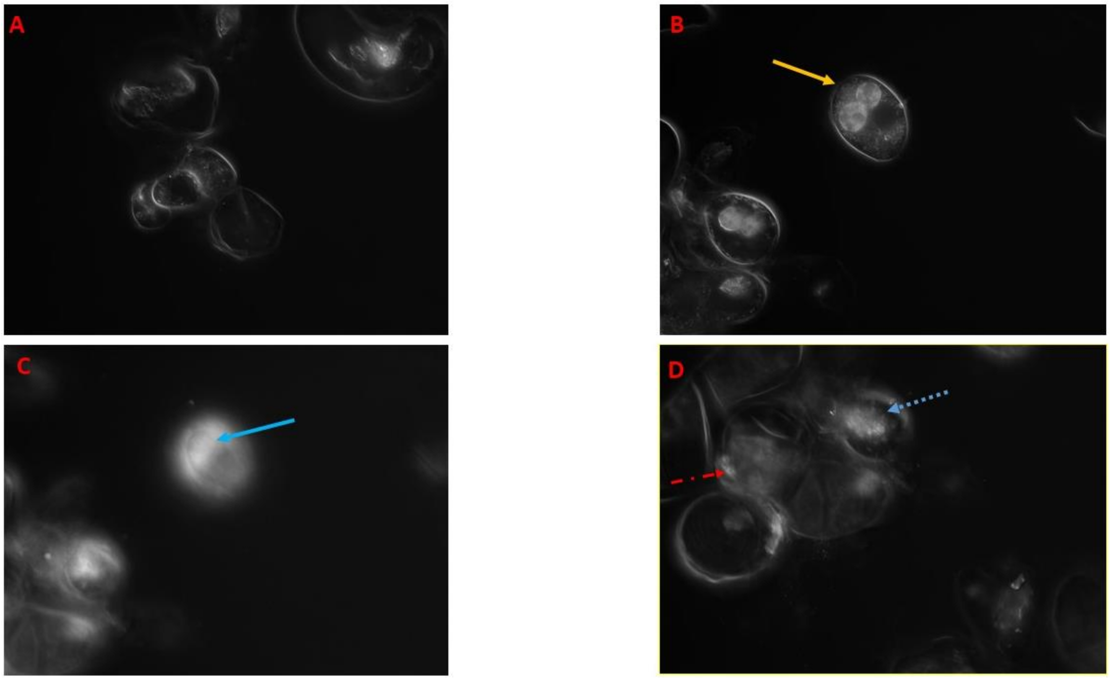


Suppl Movie 1.
Active secondary zoospores swimming inside *Pb* plasmodium in axenic cell culture. Movie clip A shows cytoplasmic stream in a healthy callus from roots of *Brassica napus*. Movie clip B is a time-lapse of the movement of secondary zoospores (yellow arrow) swimming inside infected cells in the center Movie clip. C is a Z stack of infected cells containing secondary zoospores discharged from zoosporangia, and a few multinucleated plasmodia (blue arrow). Movie clip D is a Z stack of an internal mould containing zoosporangia and islands of vacuoles (dashed blue arrow) inside the infected cytoplasm.

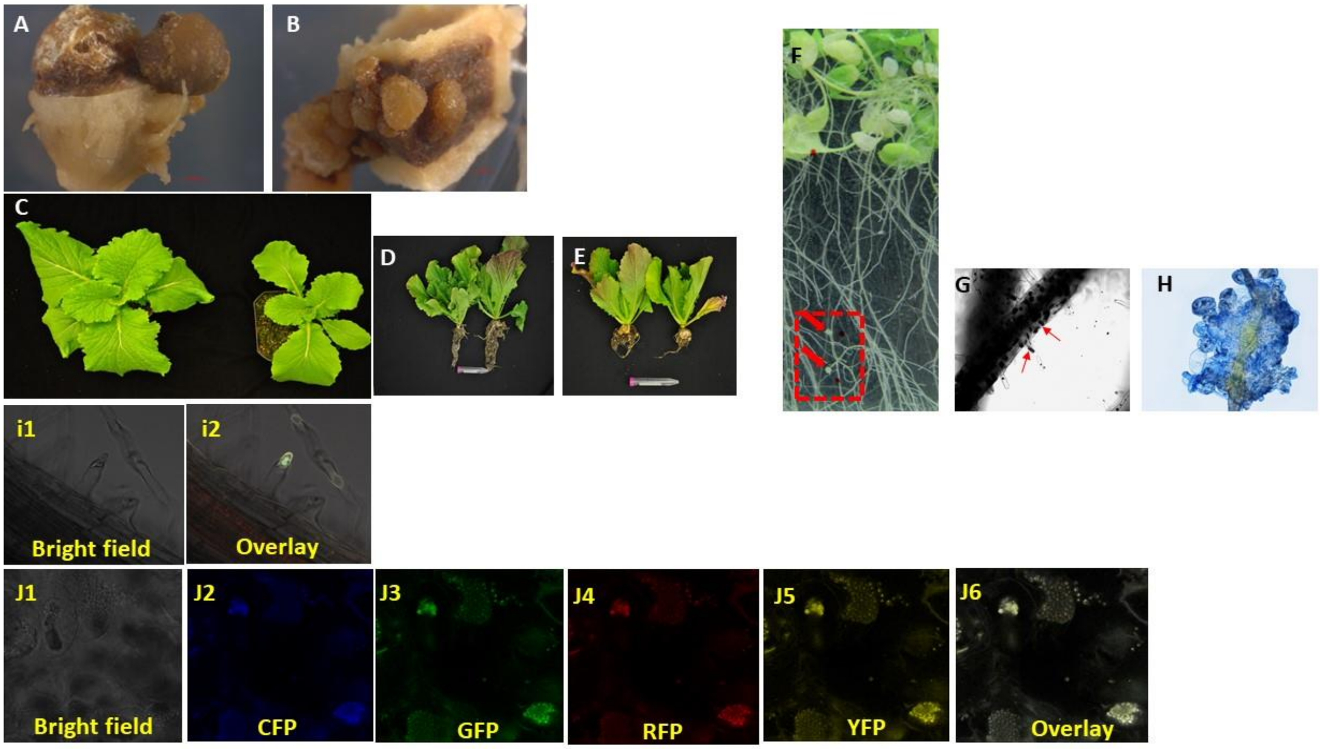


Suppl Figure 1.
Callus production and infectivity of the axenic cultures on *Brassica* and *Arabidopsis* plants. Generation of infectious callus from roots of healthy (A) or *Pb3*-infected (B) plants. Calli are generated from - around vascular bundle cells and inner cortex.
C to E is the phenotype of the healthy compared to infected *Brassica rapa* ssp. *pekinensis*cv. Granaat with RS that has been produced from axenic cell cultures of *Pb3*. C is the top view of healthy (left) and *Pb3*-infected (right). D and E show the roots of healthy and infected plants, respectively. F, G and H show infection of the Arabidopsis roots with RS that has been produced from axenic cell cultures of *Pb3*. Arabidopsis plants have been grown on the MS media and were inoculated with *Pb3* spores. F shows small galls that appeared on the roots on week three after inoculation. G is a dark-field image of Arabidopsis roots stained with trypan-blue. Plasmodial stage is observed clearly in the root hairs and cortical cells of infected plants. H projects proliferation-initiation of the gall formation on the roots of Arabidopsis. Clubbed Arabidopsis roots in section F were slightly stained with trypan-blue are shown here. i1, and i2 are pictures of a root hair of Arabidopsis infected with primary plasmodium of Pb. Pictures have been taken by epifluorescent microscope using bright field and GFP filter, respectively. *Pb* primary plasmodium is seen in green at the tip of the root hair. Pictures j1 to j5 show resting spore (RS) accumulation in the cv. Granaat roots after being infected with Pb inoculum that was increased using axenic cell cultures. J1-j6 were produced by bright field, CFP, GFP, RFP, and YFP filters of an epifluorescent microscope. j6 is an overlay picture when all the filters were applied. Figures I, and J show autofluorescence of the *Pb* zoospores and RS cells, respectively. The autofluorescence is likely due to the presence of chitin in these cells.

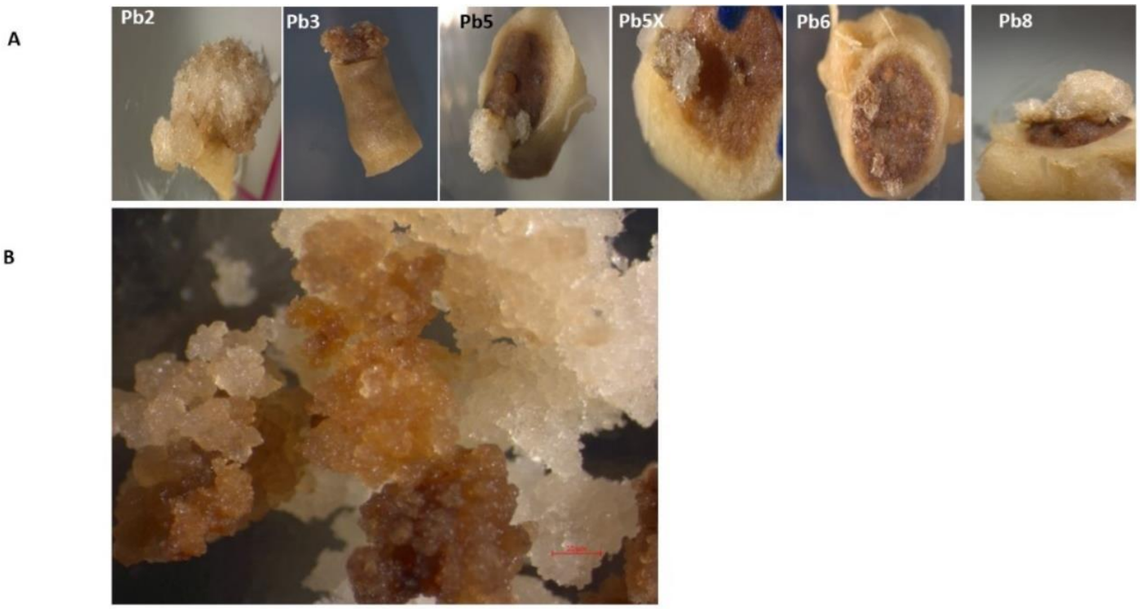


Suppl Figure 2.
Calli generated from the root segments of cv. Granaat. A) Single spore isolates of all the previously characterised Pb pathotypes based on host differentials, i.e., *Pb2*, *Pb3*, *Pb5*, *Pb5X*, *Pb6* and *Pb8* were used to infect the roots of cv. Granaat. B) Axenic cultures of Pb pathotypes in cv. Granaat produced similar calli shown in Fig. 1. All the pathotypes were able to reproduce younger plant calli in cv. Granaat transitioning from White fluffy to Brown and eventually black calli on MS media.

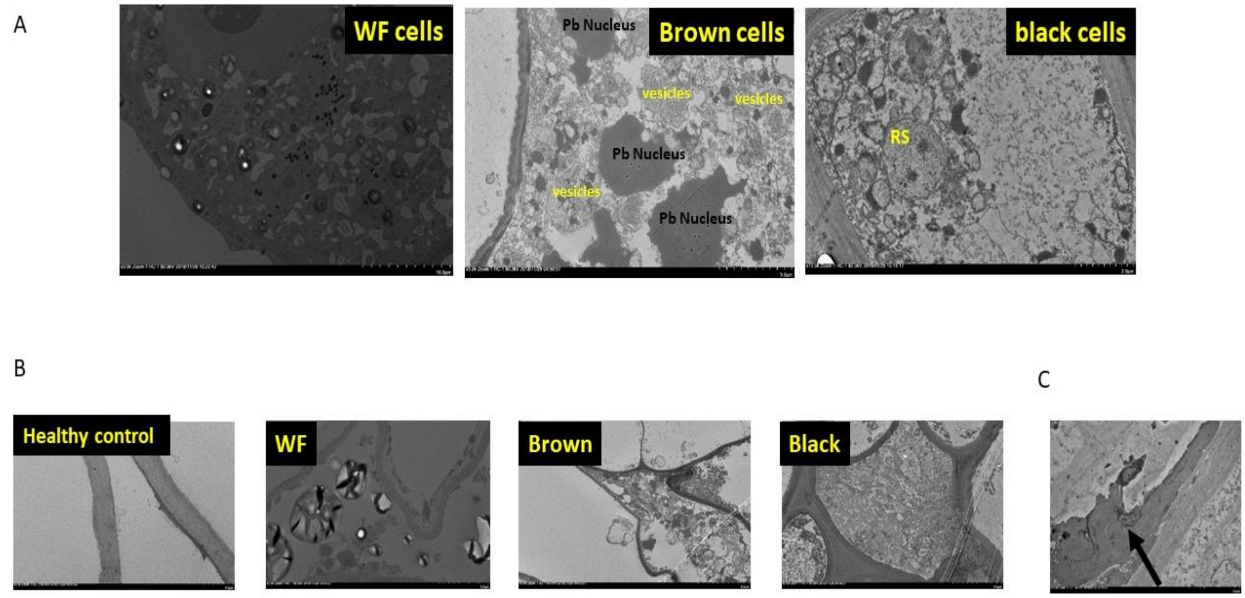


Suppl Figure 3.
Main features of *Pb* in various cells of axenic cultures. A) WF cells: Extreme induction of the biogenesis of vacuoles and induction of vesicle trafficking inside infected cells, as well as nuclear size enlargement. Various infectious stages of *Pb* including amoebae, and zoospores are observed. scale bar 20 µm. Brown cells: young plasmodia (trophozoite-like structures) surrounded by dense vesicles generated from plant cells. Scale bar 5um. Black cells: Mature RS inside black cells, scale bar 2 µm. B) Cell wall thickness. The thickness of the cell wall increases from WF to Black cells, and in comparison, with healthy control cells. Note that healthy cells only contain a large central vacuole compared to vacuolated *Pb*-infected cells. Scale bars for all pictures are 5 µm. C) Irreversible lignin deposition (Black arrow) in the infected cells adjacent to the place that pathogen is residing inside the infected cell.

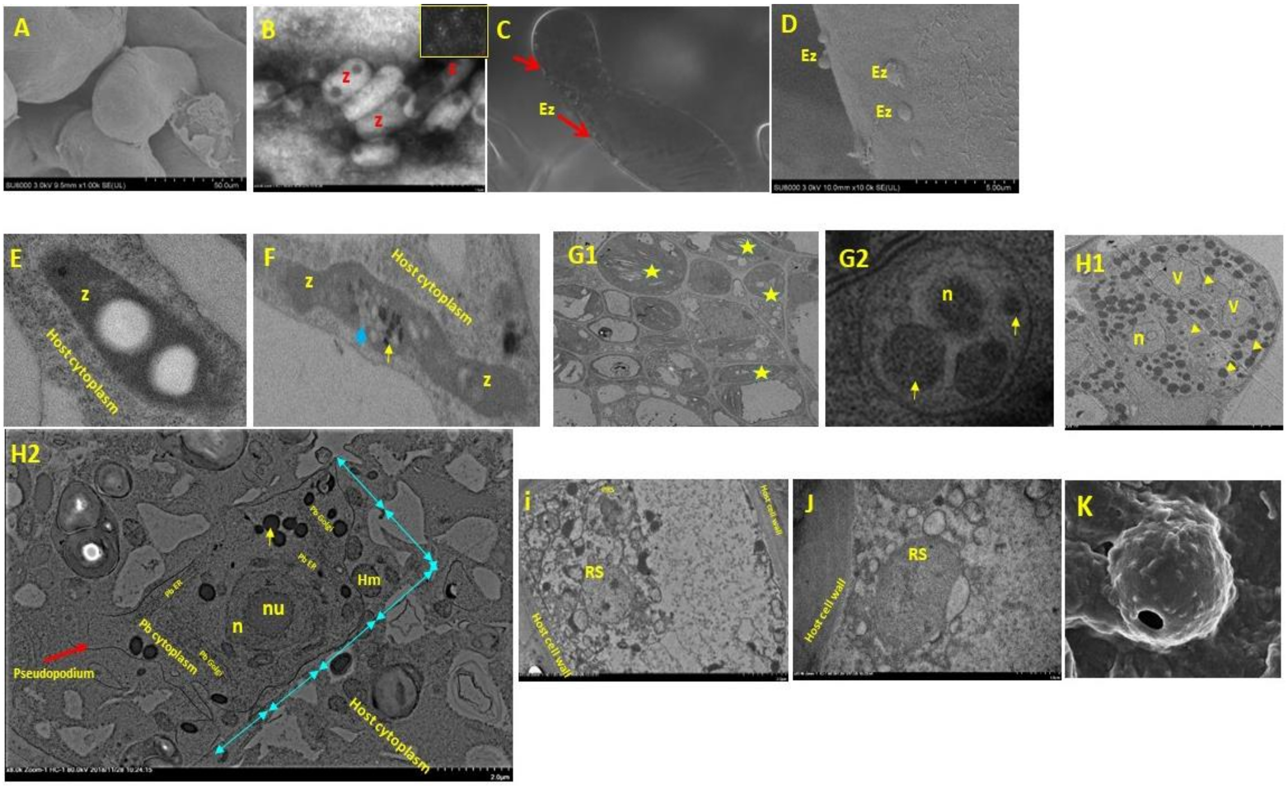


Suppl Figure 4.

Different stages of the growth of *Pb* were observed in axenic cultures. *Pb* is a unicellular cytoplasmic protist -sporogenic spore- and it changes its phenotype from one to another during infection. A-D represents external structures of *Pb*– including the production of zoospores, and their attachment to brassica cells compared to healthy cells. A) SEM of the surface of the healthy cells of *Brassica napus* cv. DH12075; scale bar is 50 µm B) Negative staining and Scanning electron microscope observation of freshly released zoospores from resting spores (RS) of *Pb*. RS cells were stored in sterile water, stained, and observed using SEM. The small square box on the right top corner shows a mass of zoospores produced. Zoospores (Z) have one or two dots at each end; scale bar is 1µm. C) LM picture of encysted zoospores on an individual somatic callus cell of DH12075. Red arrows show encysted zoospores that are attached to the cells. D) SEM picture of encysted zoospores on the cell wall of a DH12075 cell; scale bar is 5 µm. E-J represents TEM micrographs of *Pb* internal structures in infected axenic cells. E) One zoospore containing two lipid droplets within the cytoplasm of a *Pb* axenic cell culture. The black dot at one end of the zoospore is where the flagellum attaches to the cell. The size of the zoospores varies between 1-2 µm. F) Plasmogamy of two *Pb* zoospores inside the cytoplasm of an infected cell. Plasmogamy was also observed outside of the cells, when two secondary zoospores were attached (not shown). G1) Zoosporangia (yellow asterisks) containing zoospores and discharged zoosporangia. G2) Multinucleated amoeba/ primary plasmodia. H1) Premature secondary plasmodium. The transitional plasmodium -between replication and feeding stages- shows an accumulation of nuclear contents (yellow arrowhead) as well as lipid bodies like that *in Membranosorus heterantherae*. H2) Mature Plasmodium: Plasmodium has been developed into a cell which contains most of its own organelles, including Golgi, Nucleus, Endoplasmic reticulum, mitochondria, digestive vacuoles etc. Size of the amoeba at this stage is between 4 X 8 µm. The square with a dashed line is the hub of direct interaction between *Pb* and infected cells through ER. The scale bar is equal to 2 µm. i) Resting spores containing multilayer cell walls inside (RS) the black calli cells with two spots for zoospore discharge. In this picture, a premature resting spore (PRS) could also be observed. Scale bar is 2.0 µm. J) Germinating RS inside the host cell lumen. The scale bar is 1.0 µm. K) SEM of a discharged RS with a hole on the surface. n: *Pb* nucleus; nu: *Pb* nucleolus; Z: zoospore; Ez: Encysted zoospore; v: *Pb* feeding vacuole; Hm: Host mitochondria.

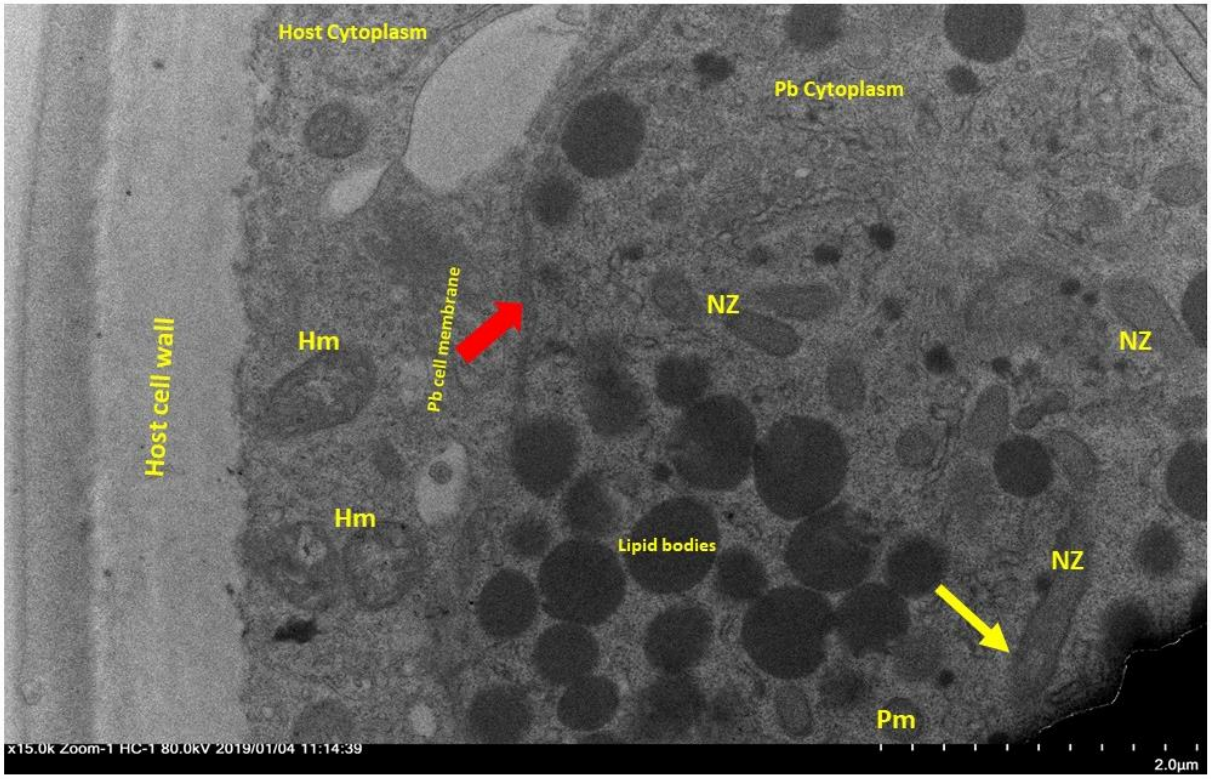


Suppl Figure 5.
TEM micrograph of a primary *Pb* plasmodium which contains secondary zoospores (NZ) harbouring retracted needle-like satchels (yellow arrow), lipid bodies within the Pb cytoplasm; Scale bar is 2.0 um. Hm: host mitochondria, Pm: Pb mitochondria.

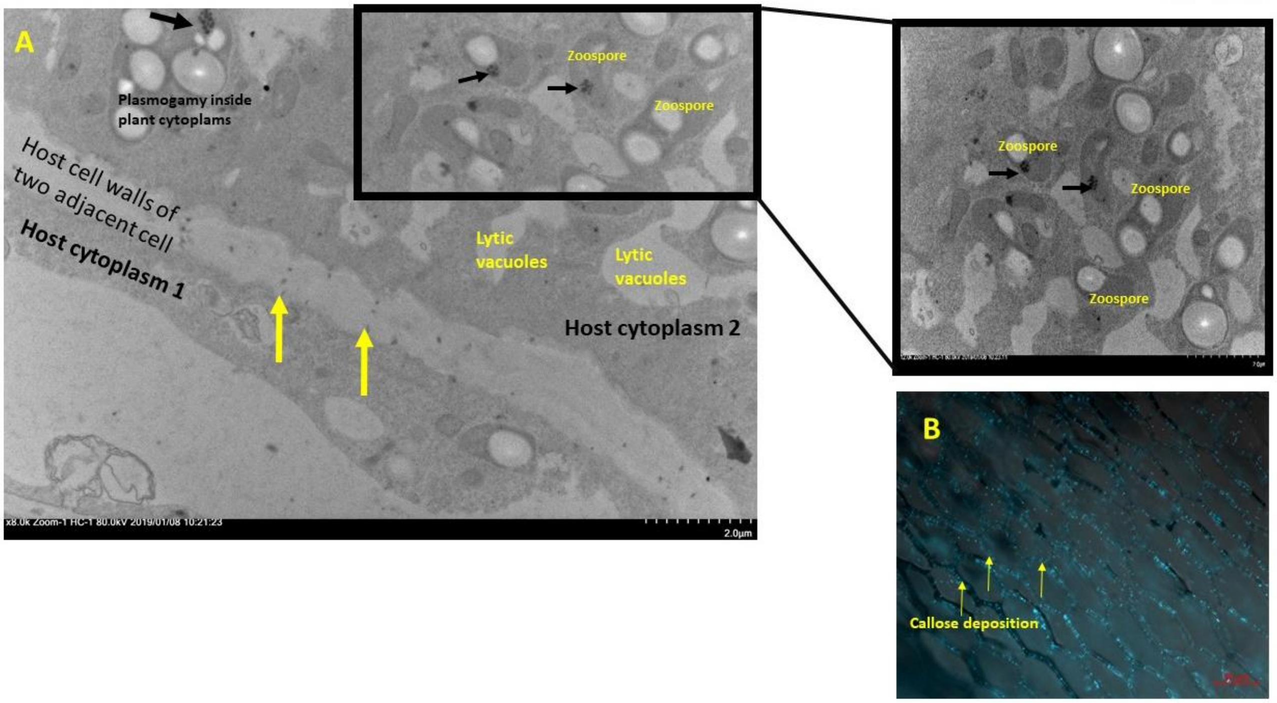


Suppl Figure 6.

Callose deposition in the host cell wall at the pathway of the zoospores between two infected cells. A) The cells were infected with *Pb* zoospores for over 12 hrs. Zoospores contain one or two oil bodies and sectional patterns of flagellar attachment to the spore- units of 9 centrioles form the core of a flagellum on one side. Axonemes (black arrows) are found in the middle/crescent side of the zoospores, but not at the basal end. The black square box shows the enlarged configuration of the zoospore cells. Zoospores are swimming between host's newly generated lytic vacuoles and defence-related vesicles to avoid them. Ineffective sequential callose deposition (yellow arrow) in the host cell wall (yellow arrows) as well as constraints in the cell wall and cell membrane is abundant. The scale bar is 2µm. A) Plasmogamy inside plant cytoplasm as well as accumulation of the secondary zoospores in the cytoplasm of adjacent host cells are notable. B) Callose deposition on the cell wall of cv. Granaat was observed upon infection by secondary zoospores of *Pb*. Roots were stained with Aniline blue and observed using epifluorescent microscope. Scale bar is 50 µm.

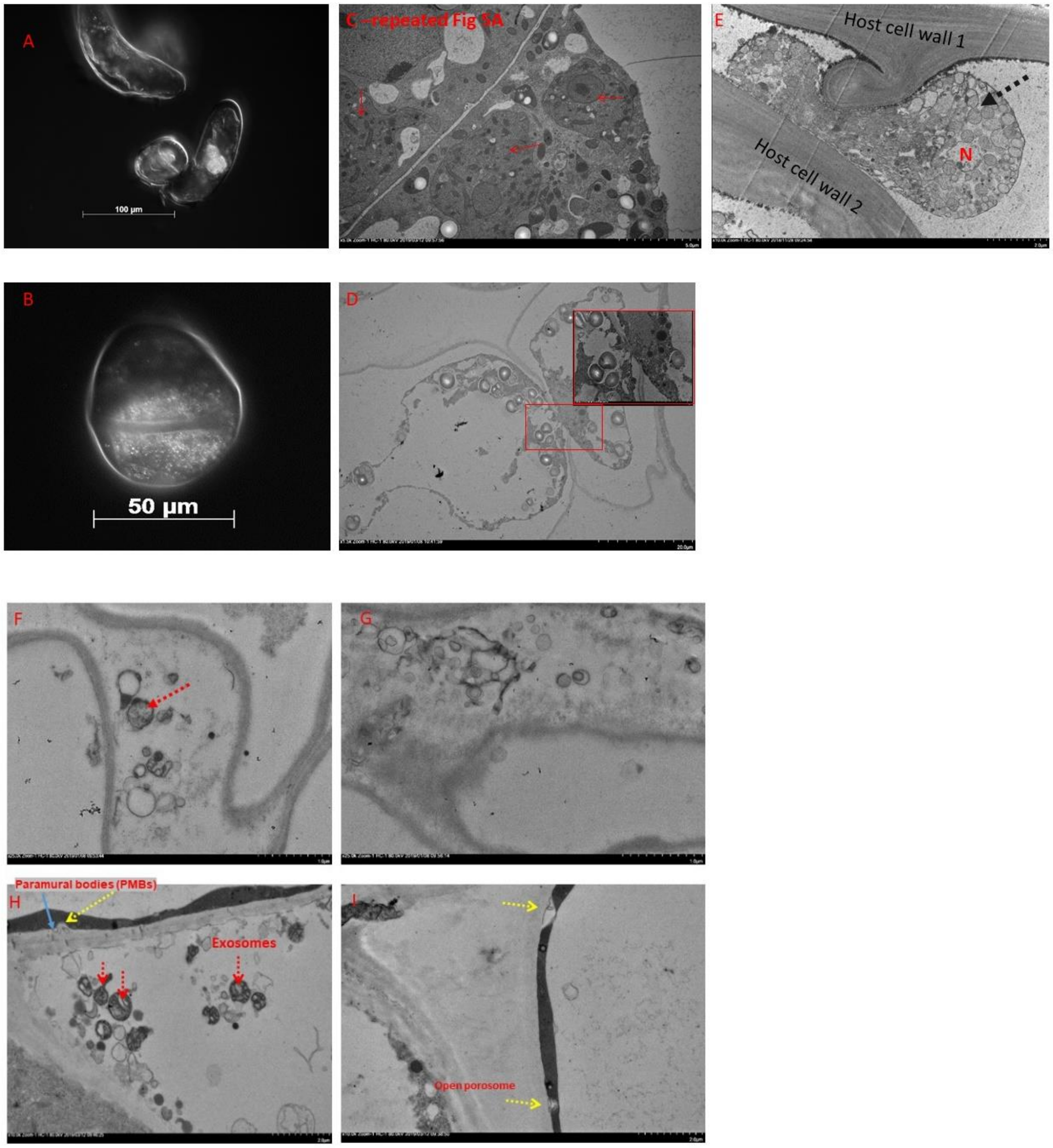


Suppl Figure 7.

Movement of *Pb* or the induced organelles in the infected cell.
A) Mitosis of the host cancerous cell while the plasmodia (A) or zoospores (B) are present in the cytoplasm. Pathogen is carried over to new cells when the host cells go through regular mitosis/cell division. Pb infection causes extensive plant cell division and enlargement. Cell division could happen at any time of the pathogen development consisting of new plasmodia, zoospores etc!
C) A TEM picture showing the separated plasmodia upon host cell mitosis. Plasmodia are shown by red arrows. Note that this picture has been presented above in Figure 5 as well.

D) The cytoplasm of two neighbouring infected WF cells that are interconnected through plasmodesmata (PD). This seems to be an efficient way of *Pb* movement between the cells.
E) Movement of a mature plasmodium between the cell walls of two black infected cells with the disintegrated plasma membrane. The mature amoeba moves in apoplastic space between neighbouring cells to locate and enter a live cell.
F) Infected Brown cells containing pathogen cells similar to the ring form stage of a malaria pathogen *Plasmodium falciparum*.
G) Accumulation of small vesicles in the paramural space of the infected black cell.
H and I) shows vacuolar accumulation of exosome in infected cells near porosomes on the cell surface. Porosomes are cup-shaped structures in the cell membranes of eukaryotic cells where secretory vesicles transiently dock and being secreted. Blue arrow in H shows the paramural bodies between PM and the cell wall that are secreted through a porosome. Red arrows show exosomes in the vicinity of porosomes.
